# Jumping over Baselines with New Methods to Predict Activation Maps from Resting-state fMRI

**DOI:** 10.1101/2020.12.15.417675

**Authors:** Eric Lacosse, Klaus Scheffler, Gabriele Lohmann, Georg Martius

**Author notes:** these authors contributed equally to this work (Gabriele Lohmann, Georg Martius).

## Abstract

Cognitive fMRI research primarily relies on task-averaged responses over many subjects to describe general principles of brain function. Nonetheless, there exists a large variability between subjects that is also reflected in spontaneous brain activity as measured by resting state fMRI (rsfMRI). Leveraging this fact, several recent studies have therefore aimed at predicting task activation from rsfMRI using various machine learning methods within a growing literature on ‘connectome fingerprinting.’ In reviewing these results, we found lack of an evaluation against robust baselines that reliably supports a novelty of predictions for this task. On closer examination to reported methods, we found most underperform against trivial baseline model performances based on massive group averaging when whole-cortex prediction is considered. Here we present a modification to published methods that remedies this problem to large extent. Our proposed modification is based on a single-vertex approach that replaces commonly used brain parcellations. We further provide a summary of this model evaluation by characterizing empirical properties of where prediction for this task appears possible, explaining why some predictions largely fail for certain targets. Finally, with these empirical observations we investigate whether individual prediction scores explain individual behavioral differences in a task.

## 1 Introduction

Functional magnetic resonance imaging (fMRI) offers noninvasive whole-brain activity measurement. Generally, different experimental paradigms are used to understand aspects of brain function. The two main experimental fMRI paradigms study the brain in *resting-state* (*rsfMRI*) and while performing a controlled *task* (*tfMRI*). The first records brain activity usually with instruction to “keep awake,” “do not think about anything in particular,” and/or “visually fixate upon a crosshair display.” In contrast, tfMRI measures brain activity evoked by tasks typically seeking to isolate some specific cognitive process, usually contrasting it to a control condition. These two paradigms are usually treated separately and little is known about how they precisely relate. However, it was observed that brain activity in both share many features that may help to explain brain function^1–9^. Many of these observations show that much of the estimated variance in rsfMRI functional connectivity (FC) appears to be shared with tfMRI activation maps. These observations are often based on group averages. However, averaging across groups destroys relevant information^10^. Therefore, predictions about individual brains are vital for making progress in neuroscience. The relationship between rsfMRI and tfMRI for individual subject prediction can be captured by a regression problem, as illustrated in figure 1. This topic has been addressed in numerous studies^11–18^. Here we re-examined methods that address this problem using machine learning techniques with only functional data. That is, learning statistical models mapping rsfMRI and tfMRI data that generalize on unseen test data (individual subjects)^19^. Problematically, when considering individual predictions evaluated over the *whole-cortex*, our benchmark comparison shows that previous methods are extremely limited beyond predicting better than a trivial baseline of group averaging. This is alarming. In this paper we develop a modification of previous methods that allows them to jump over baselines in many cases, though some limitations still exist. These modifications can be briefly summarized as follows: using a regularized regression method that fits and estimates hyperparameters on a *single* vertex or voxel basis. This technique is known from previous fMRI studies^20^, however, has not been used in this context.

**Figure 1.**
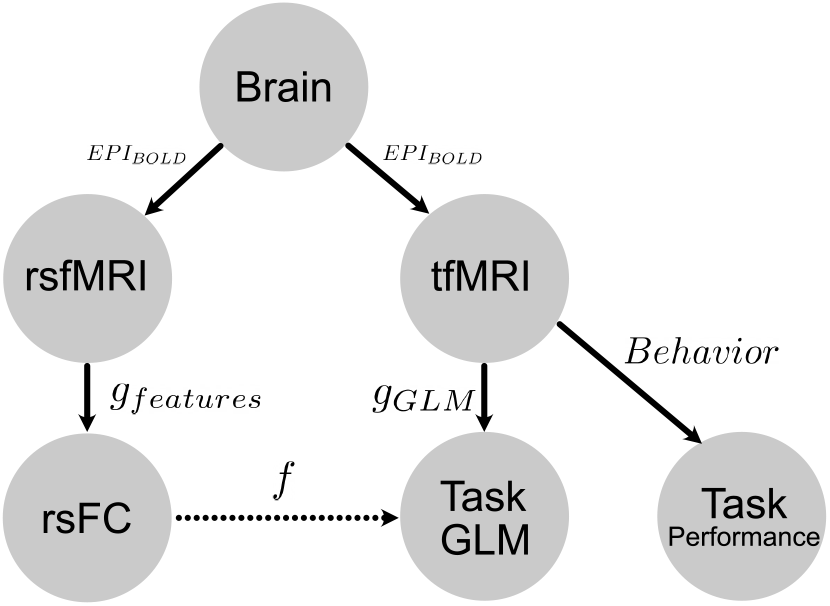
A conceptual model of the problem setup and goal. Both rsfMRI and tfMRI measuresments are acquired using BOLD echo planar imaging (*EPI_BOLD_*) by the Human Connectome Project scanners and acquisition protocols. These reconstructed and processed images from a single subject are mapped by some encoding model: either resting-state functional connectivity features (rsFC) by function *g_features_* or tfMRI data mapped to a z-statistical map summarizing task activation by function *g_GLM_*. Function *f* is the model mapping rsfMRI features to task maps. Our goal is to find optimal models *g_features_* and *f* that give the highest performing whole-cortex prediction of task GLM maps (See evaluation section for metrics describing how model performance score is measured). Additionally, activity during the acquisition of tfMRI generates some observed behavior commited during the task. Whether the relative dependence between rsfMRI and tfMRI tell us anything about task behavior is an important question we sought to answer through improving these models.

Therefore, the first aim of the present paper is to demonstrate that the methods we propose are capable of superior prediction. To do so, we provide a benchmark comparison showing how our modifications improve models considerably on a large Human Connectome Project (HCP) dataset. Following these modifications, model predictions achieve above baseline performance for a large number of target contrasts. Notably, these results not only predict individual subject differences, i.e., ‘connectome-fingerprints’^21,22^, as many have previously shown^12–18,23^; they provide support that whole-cortex prediction by a model exceeds what any kind of group averaging, i.e., baselines, could achieve–a point we will reiterate the importance of.

Second, to investigate the benefits of the proposed vertex-wise regression, we consider a set of algorithms for feature extraction and prediction, see table 1. Besides comparing relevant methods in the literature, we also provide additional insights into which features are actually predictive and discuss other aspects worth investigating. For instance, we give evidence for the relevance of the vertex-wise regularization strategy. Also, we found that widely adopted parcellations surprisingly do not outperform random projections by a considerable margin initially expected for this task.

**Table 1.** An overview of all methods we compare and benchmark. The names are composed of parts for feature extraction (**MMP**, **PCR**, **GPR**, **GICA**, **AF**) and regression model (RR, OLS), see Methods for details.

| Model Name | Proposed here | Parcellation - Feature Extraction | Type of Fitting | # of features |
| --- | --- | --- | --- | --- |
| MMP-RR-PCR | ✓ | MMP - Partial Correlations | SV Ridge Regression | 379 |
| Rest-Task GICA RR | ✓ | ICA on Task Data | SV Ridge Regression | 80 |
| Rest-Rest GICA RR | ✓ | ICA on Rest Data | SV Ridge Regression | 80 |
| MMP-RR-DR | ✓ | MMP w/ Dual Regression | SV Ridge Regression | 379 |
| MMP-RR | ✓ | MMP | SV Ridge Regression | 379 |
| GPR-RR | ✓ | Random Projection | SV Ridge Regression | 379 |
| AF-Mod | ✓ | Mean Activation Maps | SV Linear Regression | 1 |
| GICA-DR-OLS<sup>12</sup> | ✗ | ICA w/ Dual Regression | Parcel-wise Linear Regression | 50 |
| MMP-ParcelRR<sup>23</sup> | ✗ | MMP | Parcel-wise Ridge Regression | 360 |
| MMP-OLS | ✓ | MMP | SV Linear Regression | 379 |
| AF<sup>13</sup> | ✗ | Mean Activation Maps | None | 0 |

To arrive at these insights we report additional metrics that we believe should be included in these kinds of studies in the future. That is, in addition to a widely accepted metric evaluating whole-cortex predictions, we report predictive variance explained (*R*^2^ according to sum of squares) on a single vertex level. This examination allowed us to empirically investigate where predictions performed well spatially, explaining why predictions of only a certain number of contrasts perform by a respectable margin above naive baselines.

Finally, recent literature finds correspondence between rest and task activity to be rich in information about individual subject behavior^24^. Following this line, we explore the behavioral relevance of the rest-task dependency found by our best performing method. Namely, we check whether the prediction scores for individual subjects based in rsfMRI carry any information about their behavior during the tfMRI acquisition. We demonstrate how a model’s prediction score can be taken as a relative measure of dependency between rest and task measurements. In this way we show that this model may provide information relevant within a behavioural neuroscience context. We also evaluate these behavioral measures relative to a group average baseline. Our results show a compelling behavioral correspondence between resting state and a subject’s task performance in certain contrasts. We believe this can drive further progress in the field.

## 2 Materials and methods

We consider fMRI data in “grayordinate” space, an HCP-specific standard in a CIFTI data structure separating a surface cortical space that is vertex-based from subcortical and cerebellar areas that are volumetric or voxel-based. In this study we use data from the Human Connectome Project (HCP) S900 release^25^ and use 100 subjects for training and 100 subjects to make predictions. Here, we consider prediction targets of each subject *i* to be fixed-effects task GLM maps only on the cortical surface 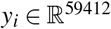 defined across 7 different task categories. Performance across these 7 task categories intended to elicit brain activity diverse enough to provide a vast coverage across the entire cortex^26^. Together, a total of 47 different contrasts were included. To model these predictions, we consider methods that first rely on some feature extraction from rsfMRI data. This feature extraction makes use of the *entire* grayordinate space, i.e., including the volumetric data component. For each subject *i* we consider the data matrix 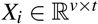 where *v* = 91282 is the vertex+voxel dimension in grayordinate space and *t* is the number of samples acquired in time. Further details on the pre-processing of *X_i_* and computation of *y_i_* are found in section 2.3.

### 2.1 Evaluation

Before detailing feature extraction and a new modeling approach, we would like to bring attention to important details regarding how the models are compared against each other. All model evaluation measuring predictive performance is done only on the 59412 cortical surface vertices within the 100 subject test-set although rsfMRI feature extraction uses the entire grayordinate space. We exclude volumetric evaluation, i.e., in subcortical regions, primarily to avoid evaluation bias due to the low signal-to-noise ratios and technical challenges of subcortical imaging^27^, for visualization purposes, and to be consistent with previous work, e.g.,^12^. Individual subject scores were computed as the Pearson correlation score *r_i_* for subject *i* between prediction **y**_*i*_ and “true” activation map **y**_*i*_. This image similarity metric is a unit-less measures that provides a concise summary of whether the overall shape of activation prediction is determined to be accurate^28^.

This measure alone, however, does not inform us *where* spatially the model is capable of making accurate predictions. For that, we include the predictive *R*^2^ score, a standard measure to quantify how much variance is explained by the predictive model^29^. Given test-set predictions at vertex *j* as 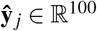 and “true” activation map 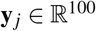, this score is computed as

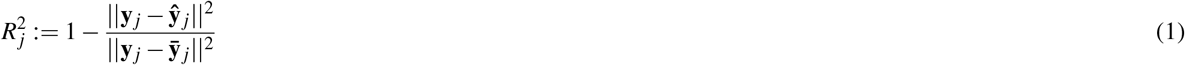

where 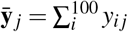 is the subject-wise mean over 100 test-subjects. This score indicates where and to what extent prediction was possible for each vertex of the fitted model. It does so by allowing a standardized comparison, i.e. as an expression of fraction of variance explained. Note that according to the definition of *R*^2^ here, it allows for negative scores. In that case, the mean of the data being evaluated would provide a better prediction than the fitted model’s output. In other words, where predictions yield a negative *R*^2^ score, predictions do not jump above a simple mean average prediction. For whole-cortex comparisons, the weighted average *R*^2^ across the cortical surface was computed. This was done by weighting each vertex *R*^2^ by the variance of the target sample.

Note that we do not report higher intra-subject vs. inter-subject prediction scores as an evaluation criterion as was done in^12^. We do not believe this observation is particularly constructive beyond the two evaluation metrics above we use. This position is based on the following observations. We understand intra-subject dependence between separate, spatially normalized whole-brain measurements exists to the extent it allows highly accurate subject identification from both rsfMRI and tfMRI-based measurements^22,30^. We could expect that an output derived from an arbitrary encoding model of rsfMRI compared to tfMRI activation maps could reveal higher intra-subject correlation than inter-subject, preserving the dependency structure defining rsfMRI and tfMRI are both acquired from the same individual brain. Yet, that prediction can be vastly poorer than a naive, unfitted baseline model in terms of whole-cortex evaluation. Supplementary figure S1 illustrates that an arbitrary FC encoding of rsfMRI can demonstrate exactly this. A correlation map produced by a random averaging can show higher intra-subject than inter-subject scores to task activation maps clearly marked. This illustrates that inter-subject differences exist despite explaining no variance on a vertex-wise level and vastly underperforming baseline scores. While this observation still reveals individual features unique to the subject are preserved, we hesitate to claim it is evidence of a successful prediction about something unknown. Instead, we believe it reiterates what we know from the very outset of the problem: both rsfMRI and tfMRI are measured from the same brain. Therefore, we try to place our claims of predictability by emphasizing comparison against models of massive subject averaging. Our goal is that our prediction performance exceed these simple subject averaged baselines across the whole cortex.

Also note that we specifically choose not to evaluate any model performance based on a measure of suprathreshold extent, e.g., thresholded maps and their overlap indices–Jaccard or Dice. We also do not report any qualitative comparisons based on suprathreshold extent as we believe it can be misleading. We found results based on these indices to be highly dependent on their chosen threshold, which acts as a nonlinear transform to spatial maps. Further, we also found that group results are highly dependent on the number of subjects used in a manner that is atypical of increasing sample size influence on model performances. That is, an increasing number of subjects used for **Group Z-stat** or **Group Z-stat (TFCE)** biases Dice coefficients scores downward when thresholds become conservative, e.g., from Gaussian mixture model thresholding. An empirical demonstration of these influences from chosen thresholds and number of subjects used on predictive versus group averaged models is provided in the supplements, figures S2, S3. These observations together provide the basis why using these two metrics appear inappropriate. Therefore, we do not use them to measure any model performance, which deviates from previous reports.

### 2.2 Modeling

The subsections below will detail various feature extraction methods used in the benchmark evaluation. For making predictions we are comparing a number of existing methods, as listed in table 1.

#### Vertex-wise Ridge Regression Model

We propose to use a regression model for *each* vertex *j* independently, each with its own hyperparameters. We use a ridge regression model fit over all train subjects *n* = 100 to estimate vertex *j* model coefficients *w* as

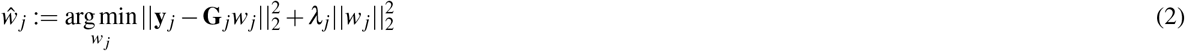

where **y**_*j*_ is an n-dimensional vector of task activation belonging to vertex *j* (here *j* = 1,…, 59412) on the cortical surface. **G**_*j*_ is the *n × f* feature matrix of extracted *f* number of rsfMRI features, as detailed below. Any model making use of this vertex-wise ridge regression is denoted by **RR** in its complete model title. Since the regression model is typically under-determined, regularization is essential for generalization of the model. We chose a quadratic regularization with hyperparameters *λ_j_* controlling the degree of regularization separately for each *j* vertex. The values *λ_j_* were chosen via a generalized cross-validation procedure over the training-set data^31^. We suspected that any method offering some degree of shrinkage would be suitable^15,20^.

##### 2.2.1 Baseline Models

Three baseline models were used to judge the actual prediction performance of all models lists in table 1. A first and most obvious choice is simply the mean (**Group Mean**) of our targets computed from the training set data. Further, we computed group-level Z-statistics with multiple comparison correction (**Group Z-Stat (TFCE)**) and without multiple comparison correction (**Group Z-Stat**) for every contrast. **Group Z-Stat (TFCE)** results were only used to investigate the results based on suprathreshold extent. Details on the computation of **Group Z-Stat (TFCE)** and **Group Z-Stat** is provided in the supplemental material.

Finally, as an additional baseline model, we fit a ridge regression model separately for each surface vertex with 6 anatomical features (**Anatomical RR**). The motivation for including the anatomical baseline stems from speculation that most variance of task-activation shapes can be explained by the subject’s anatomical features. These anatomical features are the mean image across the RL-phase and LR-phase (encoded EPI resting-state session-1 runs) and 4 anatomical T1w features extracted from Freesurfer segmentations (recon-all): cortical (quasi) myelin, sulcal depth, curvature, and thickness maps.

##### 2.2.2 Resting-state Feature Extraction

All resting-state models we consider rely on some functional covariance-based (FC) feature extraction of resting-state data of the entire grayordinate space. For each subject *i* the normalized data matrix 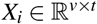 is converted into the feature matrix *G_i_* in the general form as:

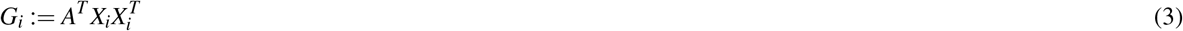

where 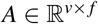. *A* projects the subject-sample covariance matrix into a lower dimensional space *f* (number of features). *G_i_* is also known as a “semi-dense connectome”. Matrix *A* is selected either based on predefined Regions of Interest (ROI), e.g. parcellations, on group-based ICA, or on selecting specific features directly, e.g., random projections or mean task activity. Note, that no smoothing of data matrix *X_i_* was applied before any feature computation in any of methods examined.

###### Multimodal Parcellations

Let us first consider the case of predefined brain regions using Multimodal Parcellations^32^ (**MMP**) with *f* = 379 and *A* ∈ {0,1}. In other words *A* averages over the activity in spatial regions. As an additional modification we include an additional step of Dual Regression for feature extraction^33^ denoted as **DR**.

###### ICA-based

For the case of ICA methods we consider the method of computing *A* via Multi-subject/Group Independent Component Analysis (GICA) for calculating *G_i_* following the algorithm (Canonical ICA) outlined in^34^. This was done to compare *group maps* extracted either between rest or task. That is, **GICA** features were either derived from rsfMRI or tfMRI data for models **Rest-Rest GICA RR** or **Rest-Task GICA RR**, respectively. In the tfMRI case, separate features were calculated by selecting only 6 of the 7 tfMRI datasets, leaving out the tfMRI measurement of the to-be predicted GLM task contrast. Doing this excludes circularity. These group-level maps were computed over the 100 training subjects. Briefly, the estimation involved a separation of subject-level noise by applying PCA in the time dimension. These subject-level PCs were then concatenated to estimate group-level patterns via Canonical Correlation Analysis (CCA). Group-level PCs were then finally decomposed into group-level independent sources with ICA via FastICA^35^. The number of both subject-level and group-level components selected was 80^36^. Note, we did not apply further region extraction from these group-level maps to obtain non-overlapping, parcellations. Hence, the number of features remains at 80.

###### Activity Flow

A method called “Activity Flow”, **AF**^13^ uses a group-mean task-activation pattern computed across the training set and uses it directly for prediction for held-out regions of the cortex (as defined by some parcellation) without data-driven fitting. Note, like all other models, we do not perform spatial smoothing on the rsfMRI data. We add a version where this is selected as a single feature used for regression called **AF-Mod**.

###### Random projections replacing parcels

To assess the impact of the parcellation, we replaced the standard parcellation with a random projection scheme. Random projections are a technique for dimensionality reduction using a random matrix having unit column norms such that the projected lower-dimensional subspace approximates the original distances between data points. Provably, if data points in a vector space are projected onto a randomly selected subspace that is sufficiently large, distances between data points are approximately preserved^37^. In our case *A* is a randomly generated matrix drawn from 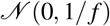 denoted as Gaussian Random Projection (**GRP**).

###### Principle Component Regression

Functional correlation features extracted from **MMP** models so far do not distinguish between direct and indirect interactions of whole-cortex brain activity to time-dependent signals averaged within parcels. In order to compute features that resemble direct interactions more closely, principle component regression **PCR** is used to compute a semi-partial covariance feature matrix *G_i_* for each subject^13,38^. This was accomplished by masking vertices for exclusion within a crucial area surrounding each parcel. Since neighboring vertices are spatially autocorrelated, this step is essential. In detail, **MMP** partial covariance matrices were computed for each subject by projecting a masked data matrix 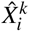 for each ROI, *k* = {1,…, 379} belonging to the **MMP** parcellation and masking all surrounding vertices within a 10mm neighborhood of vertices belonging to the *k^th^* ROI. Surface cortical distances were estimated as their geodesic distances on a group-averaged (all 200 subjects) midthickness surface mesh. Subcortical distances were estimated by their Euclidian distance within MNI space. For every masked ROI, 512 principle components (PC) were computed via a randomized singular value decomposition (SVD)^39^. These selected PC covariates were then regressed using ordinary least squares (OLS) onto the selected *k* ROI mean signal averaged time-series. Estimated regression coefficients from this regression were then projected back into the original 91282 dimension space of the original data matrix *X_i_*. This together results in a same sized subject feature matrix *G_i_* based on **MMP** as used in other models that only compute covariances.

##### 2.2.3 Modified Activity Flow Model

As mentioned above, the Activity Flow model performs no statistical fitting to task activation maps. We include our technique of vertex-wise regression to the **AF** model, denoted as **AF-mod**. In detail it is learning a simple two parameter OLS model fit of 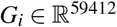 to task maps for each surface vertex. This was similar to our other vertex-wise models. Additionally, model **AF-mod** does not perform spatial masking of vertices surrounding the ’to-be’ predicted vertex as done in the original Activity Flow conceptualization. We do not perform region or vertex prediction in held-out regions.

##### 2.2.4 Remarks on Method Choices

Importantly, we *only* use BOLD data features for all resting-state data model evaluations since this is what underlies our significance claims, deviating from^12^. Also, a 100/100 train/test split was used rather than the leave-one-out cross validation employed in^12,13,15^. In all cases, all features for each subject were normalized to zero mean and unit norm. Note that we did not seek to use an optimal cross-validation strategy to maximize the performance available on the whole dataset, but provide a robust comparison of generalization performance across models given a large test sample size case. Lastly, due to the enormous computational burden of computing a vertex-wise semi-partial covariance matrix, we do not to implement the partial covariance model described in^13^. To do so would be an enormous computational burden that would require downsampling the data since a PCR would need to be computed at each vertex for each subject. Additionally, downsampling the data would render model comparison unfair between models. All evaluations of all model performances were across the same sized data with no additional smoothing applied. Further details regarding model implementation of the Group ICA dual regression OLS model **GICA-DR-OLS**, vertex-wise Activity Flow **AF**, and a ridge regression model fit over parcellations rather than single vertices/voxels **MMP-ParcelRR** may be found in^12^,^13^,^15^, respectively.

### 2.3 fMRI Data and Processing

All data analyzed in this study is from the Human Connectome Project (HCP) S900 release^25^. To limit a number of covariates that are known to be severe confounds to any of the inter-subject analyses, we selected 200 *unrelated* subjects, i.e., no family relatives, with a T1, T2, complete rsfMRI, complete tfMRI, and physiological data acquired. Additionally, we selected subjects with functional data reconstructed exclusively with algorithm r227. From these available subjects, a random selection of 100 males and 100 females were made.

The study was performed using data provided by the Human Connectome Project (HCP). All data accessed, downloaded, and used by this study was in accordance with WU-Minn HCP Consortium Open Access Data Use Terms (https://www.humanconnectome.org/study/hcp-young-adult/document/wu-minn-hcp-consortium-restricted-data-use-terms). The study was performed in agreement with those terms. By agreeing with those use terms, no further ethics approval was required at our local institute to use the data. The HCP project (http://www.humanconnectomeproject.org) is an open National Institutes of Health (NIH) initiative and received the required ethics approval for data acquisition and public distribution. All subjects who participated gave written, informed consent according to the protocol by the HCP consortium as approved by the Washington University in St. Louis Institutional Review Board (IRB). All human data was acquired in accordance with these experimental procedures adhering to these IRB processes by the HCP. These can be found in further detail^25^.

All results in this manuscript are performed on a random train-test split (100/100 subjects) of the 200 selected subjects.

Functional data was acquired with highly accelerated gradient echo type echo-planar imaging (GRE-EPI) in 2 sessions on 2 separate days with 2 two different phase encoding directions (left-right and right-left). These 4 runs, 15 minutes each, were acquired with the behavioral instruction to keep eyes open with fixation on a projected cross-hair^40^. All runs were concatenated together prior to deriving rsfMRI features. 7 tasks were performed during the task functional acquisition (IDs: emotion, language, motor, social, gambling, relational, working memory). Further details regarding the tfMRI paradigms and the extent of their brain coverage is found in^26^. Due to some of the potential benefits offered by particular HCP data acquisition choices, data used for our analyses were exclusively in the standard CIFTI-grayordinate space form. This form allows combined cortical surface and subcortical volume analyses without enormous storage and processing burdens among increases in SNR due to surface smoothing and and better cortical fold alignments^41^.

Minimally preprocessed ICA-FIX denoised data of the HCP was used for our analysis. Details and code of those pipelines can be found in^41^ and^42^, respectively. Each measurement had its first 5 repetitions discarded before any local processing. All data prior to being applied in any of the models implemented were demeaned and variance normalized (unit-noise variance) feature-wise. No additional preprocessing procedures, e.g., filtering or smoothing, were applied.

Prediction targets were fixed-effects (2 Sessions) GLM estimated contrast maps over all 7 tasks with a surface smoothing kernel FWHM of 4mm applied. Fixed-effects GLM results were computed by HCP tfMRI pipelines in CIFTI-greyordinate space and z-transformed^42^. All HCP tfMRI pre-computed GLM contrasts from these tasks are used such that no redundant predictions would be made, e.g., from sign flipping the contrast vector. This selection follows^12^ such that 47 contrast map targets are used.

A cortical parcellation with 360 regions generated by the work of^32^ was used for the left, right cortical surfaces, and we refer to this parcellation as **MMP** (MultiModal Parcellation). Additionally, for completeness and to utilize the volumetric data component of CIFTI-greyordinate space data for feature extraction, we used an additional 19 sub-cortical regions parcellation given by the HCP release, available at^42^. This results in a total of 379 regions.

### 2.4 Behavioral Data

An assessment of cognitive ability of individual subjects was provided by measures tested during tfMRI acquisition (downloaded at https://db.humanconnectome.org). This is used to understand whether individual predictions scores are related to the amount of correspondence between rest and task. Here, we correlate prediction scores (See Evaluation section) to individual behavior measures of cognitive ability. The cognitive tasks for our analysis are behavioral measurements during: working memory, language, and relational processing tasks performed while inside the scanner. Following^24^, these tasks were selected primarily because they fulfill normality assumptions. Additionally, they provide the most complete tasks associated with the contrasts we choose for predictions. Pearson R correlation prediction scores were all Fischer-z transformed across all subjects, a variance-stabilizing transformation, before computing further correlations between the behavioral measures.

#### Software Implementation and Usage

Python was used for all reported experiments and implementations with the exception of model **GICA-DR-ICA**. This model was implemented in Matlab using code shared from the authors^12^. Scikit-learn provided state-of-the-art statistical learning algorithms (http://scikitlearn.org)^43^. Additional experiments used code modified from the nilearn library for high-dimensional neuroimaging datasets (http://github.com/nilearn/nilearn)^44^. Flatmap cortical visualizations used code modified from^45^. The neuroinformatics platform that allowed downloading large datasets and a tool for 3D cortical visualizations used software provided by HCP^46^. Our public code is available at https://gitlab.com/elacosse/cf-benchmark-dev).

## 3 Results

### 3.1 Benchmarking: Which Methods Jump over the Baseline?

First, we investigate the accuracy of predictions using the described methods based on Pearson *r* correlation score for individual subject prediction. We provide a comprehensive performance benchmark comparison with a total of 14 different models. These are compared across the 47 contrast-map targets provided by the HCP S900 dataset. Note that we only focus on model prediction of a single contrast map; this does not leverage any additional information provided by incorporating multiple maps for prediction across subjects.

Our benchmark evaluation compares models using resting-state data against each other and, importantly, against simple baselines models. This is reported in two figures 2, 4, summarizing results across the entire cortex according to either Pearson *r* correlation scores or vertex predictive *R*^2^ scores (Eq. 1). The scores displayed in figure 2 are provided in the supplements table S1. All models were evaluated with the same test-set consisting of 100 subjects. This allows to report statistical significance with a one-sample paired t-test. Importantly, **Group Z-Stat** (r = 0.540 ± 0.044) shows Pearson *r* correlation score mean performance worse than **Group Mean** (r = 0.561 ± 0.047) for the vast majority of contrasts; only four (three making up the worst performing contrasts) from **Group Z-Stat** performed significantly better than **Group Mean**. Additionally, a model fit only from anatomical features **Anatomical-RR** does not generalize better than **Group Mean** baseline across all but one, the highest scoring contrast (REL). Therefore, comparisons are made against **Group Mean**, the highest performing baseline model. Many methods, especially from previous approaches, fall short of jumping over this trivial baseline, meaning whole-cortex prediction from the resting state are problematic. That is, despite many of these contrast’s Pearson correlation scores appearing quite high. However, results marked in figure 2 by significance boxes reveal that only a limited subset of the 47 contrasts do significantly better than a group mean baseline, **Group Mean**. The margin of difference between predicted score and mean baseline is shown in supplementary figure S5.

**Figure 2.**
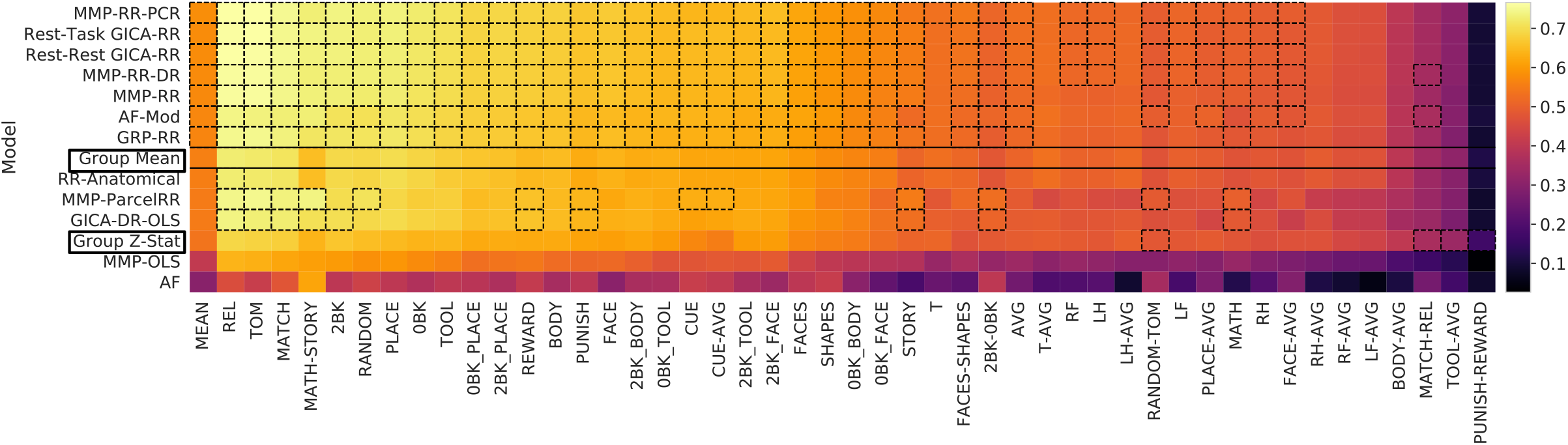
Pearson *r* correlation score benchmark results for 100 subject test set: Colorbar indicates mean *r* score across all test subjects for given contrast and model. Dashed black boxes indicate where model performance is significantly greater than test-subjects’s baseline (mean) model performance (one-sided paired sample t-test, *p* < 0.05, 5000 permutations, Bonferroni corrected across contrast comparisons). Boxes in the left column mark baseline models. Scores are ordered top (best) to bottom (worst) by their subject-wise mean score computed across all 47 different contrasts (left-most column).

#### 3.1.1 Improved predictions by vertex-wise models

All methods *with* the proposed vertex-wise fitting procedure demonstrate subject predictions (averaged across contrasts) above the mean baseline prediction (**Group Mean**), figure 2 (one-sided paired sample t-test, *p* < 0.05, Bonferroni corrected across all 47 contrasts, 5000 permutations) and figure S5.

Our model **MMP-RR-PCR** yields both the highest mean performance of subject scores averaged over all contrast targets (r = 0.582 ± 0.048) with the highest number of significant prediction performance, see figure 2. Additionally, this model holds the highest performance in 31 of the 47 contrasts (see table S1). However, several other models augmented with our vertex-wise regression method show only slightly worse performance, as figure 3 highlights. A direct comparison between the classical way of tuning the ridge regression parameter and our vertex-wise method is seen by comparing **MMP-RR** (r = 0.574 ± 0.048) versus **MMP-ParcelRR** (r = 0.550± 0.049), showing a significant gain (one-sided paired sample t-test, *p* < 0.001, t = 15.77, 5000 permutations).

**Figure 3.**
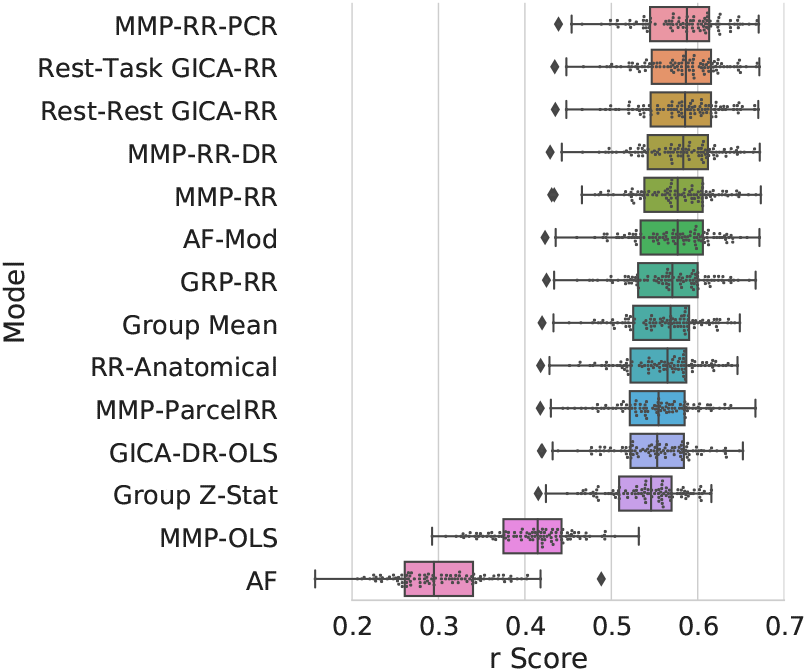
Pearson *r* correlation score results for 100 subject test set averaged across contrasts. Models are ordered top to bottom by score. These scores appear as the first column in Figure 2. Top performing models perform similarly between each other. Individual subject scores from the test-set are plotted along with box-whisker plots showing quartiles of prediction score distribution.

To understand the importance of regularization, we can compare **MMP-RR** (r = 0.574 ± 0.048) and **MMP-OLS** (r = 0.409 ± 0.047), where the latter only relies on ordinary least squares fits. This notable performance difference shows that regularization is essential for successful generalization when the number of rsfMRI features is very large. However, a complex model is not necessarily needed for successful prediction; Model **AF-mod** (r = 0.571 ± 0.049) generalizes comparatively well and has proven to be one of the best performing models despite its simplicity. From our analysis, we expect many methods with some degree of shrinkage would reveal comparable performance when trained on a single-vertex level^15,20^.

#### 3.1.2 Effects of Feature Extraction and Parcellation

We investigate the effect of various feature extraction strategies for determining *A* in eq. 3. First, *A* derived from task **Rest-Task GICA** data yields a very small improvement over model **Rest-Rest GICA** derived only from resting-state data, see table S1. This motivated us to investigate other effects of selecting *A*. Specifically, we replaced the expert-based parcellation **MMP** with a random projection *A*. Again, the advantage of a expert-based parcellation over a random projection is surprisingly small: **GPR-RR** *r =* 0.568 ± 0.048 vs. **MMP-RR** *r =* 0.574 ± 0.048. This result suggests that in many cases random projections for generating features appears to be sufficient. It simply provides a means of performing dimensionality reduction akin to perhaps any arbitrary parcellation scheme, an observation consistent with^15^.

Lastly, we investigate whether deriving more subject specific features via dual regression yielded any appreciable improvement. Model **MMP-RR-DR** over **MMP-RR** shows a statistically significant, yet small, improvement over subject predictions averaged across contrasts (one-sided paired sample t-test, *p* < 0.001). For small sample sizes, however, the use of dual regression appears to be promising, see supplementary figure S9.

### 3.2 Predictive *R*^2^ Evaluation

In addition to Pearson *r* correlation scores, we examine the variance explained on a vertex level (equation 1) evaluated on the same test set. This evaluation is summarized in figure 4 and provides a complementary measure of prediction performance. The scores displayed in figure 4 are provided in the supplements table S2. To quantify one number per contrast we report the variance-weighted average of the *R*^2^ scores across the cortical surface. This number is color-coded in figure 4 and quantifies to which degree and in which contrasts predictions about individuals can be made. Models and contrast targets with a positive *R*^2^ aligns well with the ordering of previous figure 2 results and supports how the use of single-vertex regression based methods yields a considerable performance boost and valuable predictions. Nevertheless, figure 4 emphasizes that it is only roughly half of the contrast targets that show considerable predictability.

**Figure 4.**
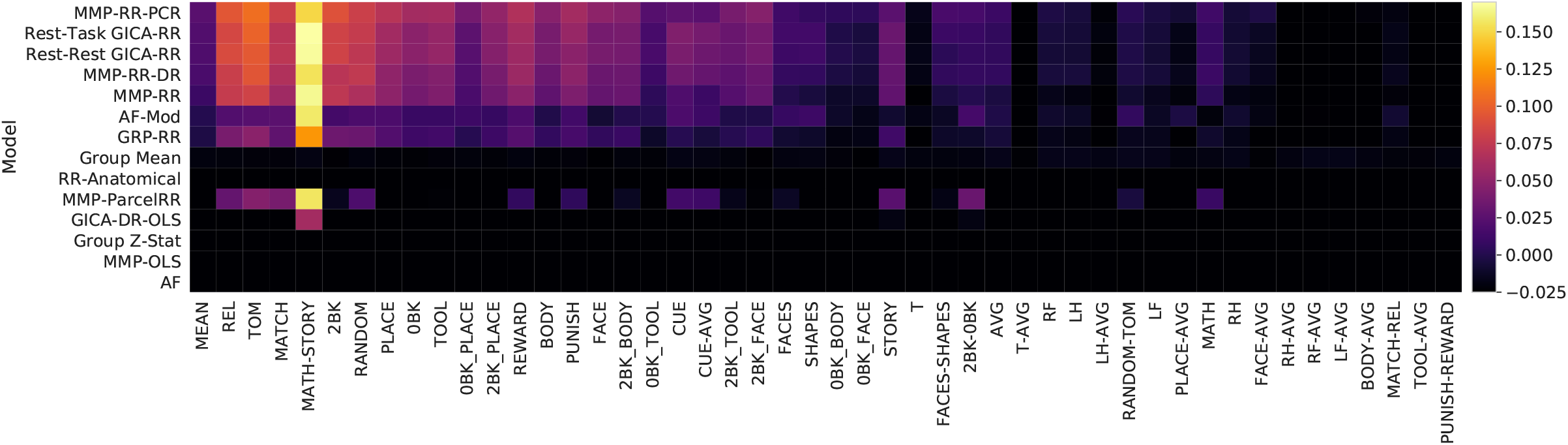
Predictive *R*^2^ score benchmark results: scores indicate the mean of cortical surface *R*^2^, see Eq. 1, weighted by the variance of each surface vertex. The colorbar indicates this measure. Roughly half of contrast targets have mean cortical *R*^2^ below 0 since predictive *R*^2^ can be arbitrarily negative. Math-Story stands out as the easiest contrast to predict. A discussion providing a reason why is provided in section *Spatially resolved predictability*. Column and row ordering are not sorted by performance and remains identical to figure 2. The left most column is the mean score across all contrasts.

#### 3.2.1 Spatially resolved predictability

Figure 4 shows considerable variability between predictive performance of certain contrasts. This can be explained due to the fact that only certain regions of the cortex drive a model’s prediction ability above the baseline. This becomes clearer with an investigation of where on the cortical surface we observe positive *R*^2^ values. To report this concisely, we render the cortical surface with a mean averaged *R*^2^ score across the 47 contrasts of model **MMP-RR-PCR** in figure 5. An additional plot showing individual contrast *R*^2^ across each task category separately is shown in figure S4. The surface plot reveals that only a limited subset of vertices lying outside of the primary-sensory regions can explain the 100 test-sample variance. These remain confined within the association cortex where most inter-subject variability of rsfMRI functional connectivity lies^47^. Regions of high inter-subject variability as measured by either rsfMRI features, task activation maps, or sulcal depth of a subject’s brain anatomy are associated with the predictability, see figure S6. This outlines that regions where subject differences in the cortical functional anatomy are highest are the regions where subject rsfMRI features or task activations also differentiate themselves the most. Supplementary figure S7 shows this spatially in flatmap visualization.

**Figure 5.**
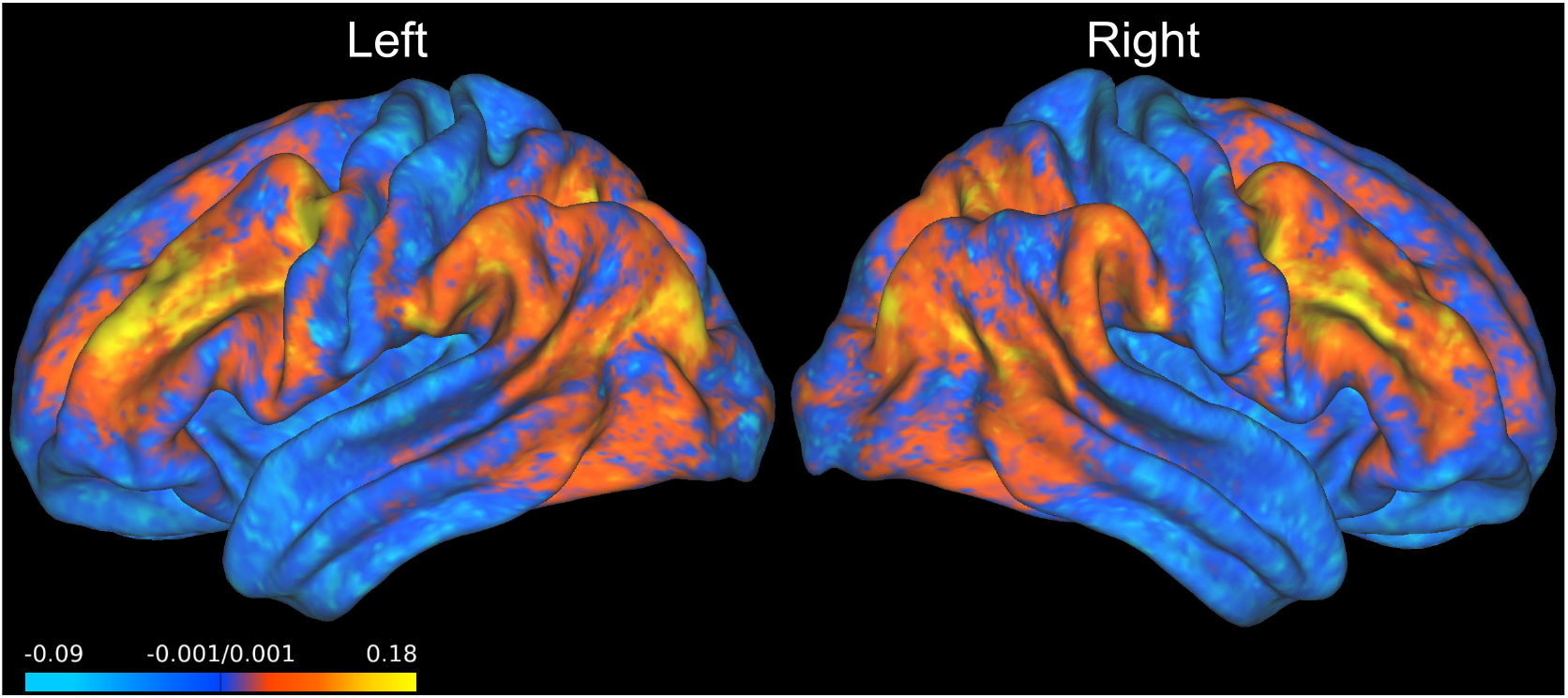
Mean *R*^2^ Score of MMP-RR-PCR across all contrast targets. Plotted are the *R*^2^ values averaged over the 100 test subjects. This is rendered on the a 200 subject averaged midthickness surface map of left and right cortical hemispheres. Positive values (red and yellow) indicate where prediction is possible. Note that prediction accuracy is best outside the primary sensory regions.

To give a better empirical characterization of the spatial dependency of model parameters and prediction quality, we report several metrics per vertex for the **MMP-RR-PCR** model. For visualization we use flatmap cortical projections of the entire cortex, as shown in figure 6. We consider the root mean square errors (RMSE) in figure 6(A) and see that the highest RMSE appears primarily concentrated around the visual cortex. The vertex-wise strength of regularization *λ* determined via cross-validation over the training-set is shown in figure 6(B). Strong regularization is employed in primary-sensory regions where predictions perform poorly. The optimal regularization is inversely proportional to the explained variance shown in figure 6(C,D). We show *R*^2^ on the training subjects (C) and on the 100 test-subjects (D).

**Figure 6.**
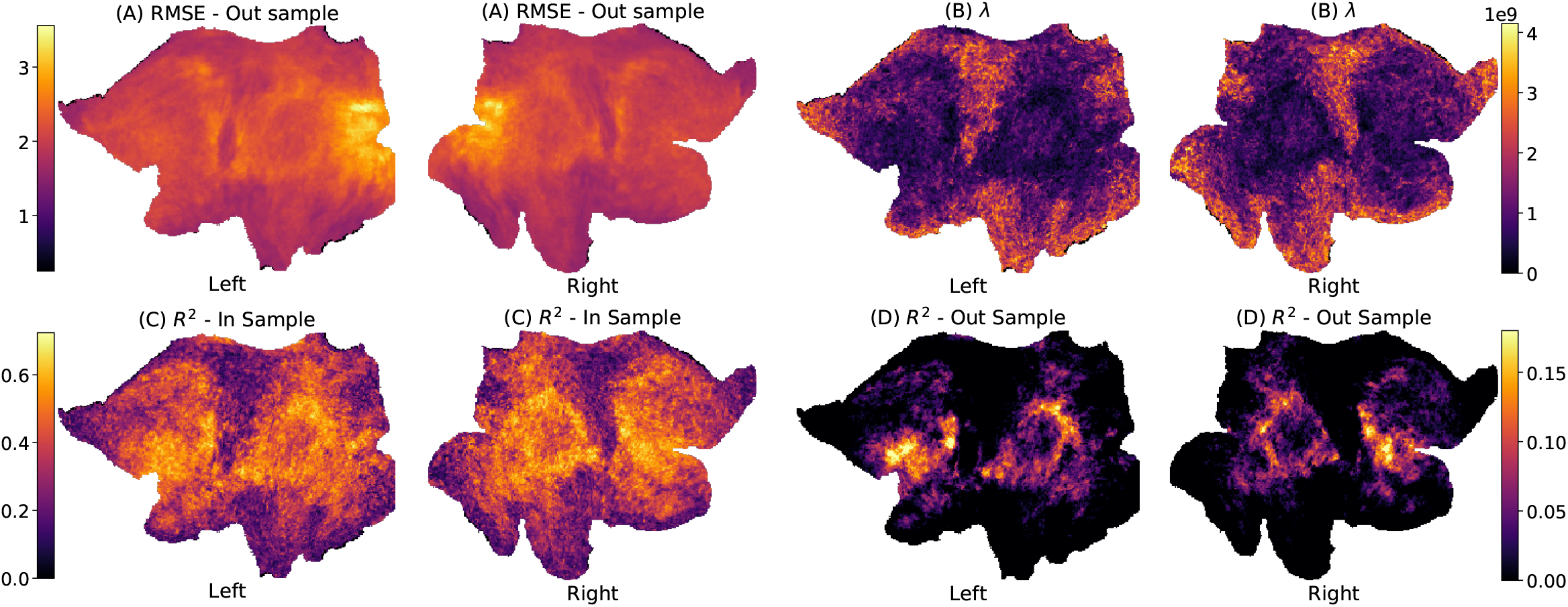
Flatmap cortical surface projections of MMP-RR-PCR model fits. (A) Root Mean Square Error (RMSE), (B) Degree of regularization *λ* in model fit, (C) *R*^2^ – 100 train subjects (D) *R*^2^ – 100 test subjects. RMSE, *λ*, and *R*^2^ are averaged across all 47 model fit results. Train and test *R*^2^ show consistent patterns between each other. *λ* shows how regularization is inversely related to the method’s ability to predict (*R*^2^). Both vertex-wise *R*^2^ and regularization parameter *λ* offer the ability to resolve spatially where rsfMRI data is capable of any prediction.

### 3.3 How many subjects are needed?

To examine top performing models closer and according to their capacity, we investigate the impact the number of training samples on 4 of the best models (**MMP-RR-PCR**, **Rest-Task GICA-RR**, **MMP-RR-DR**, **AF-Mod**) as defined by their median contrast score (left most column in 2. We included two baseline models **Mean (Baseline)**, **Group Z-stat** for comparison. These models were all evaluated on the 100 subject test set. 3 contrast targets were arbitrarily chosen because of their poor, mediocre, good performance as contrasts Motor–Right Hand, Emotion–Faces, Language–Math-Story, respectively. Pearson *r* correlation scores and predictive *R*^2^ score with respect to the number of subjects (3-100) are reported in figure 7 and figure 8, respectively.

**Figure 7.**
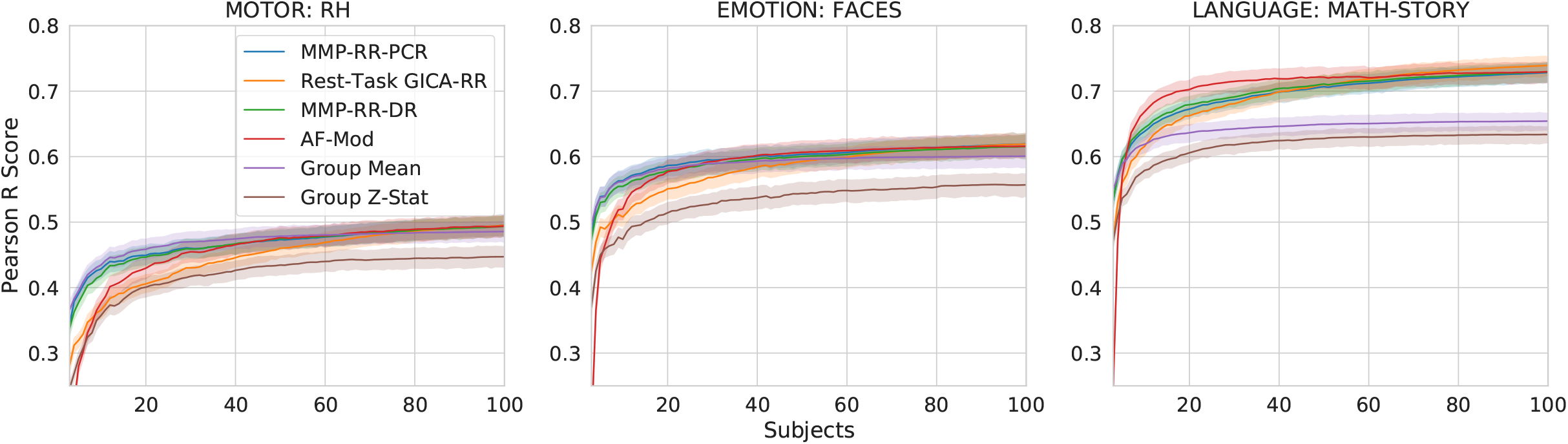
Subject-wise Pearson *r* score benchmark results for 3 selected (poor, mediocre, good), 4 high performing models (**MMP-RR-PCR**, **Rest-Task GICA-RR**, **MMP-RR-DR**, **AF-Mod**) and two baseline models (**Mean (Baseline)**, **Group Z-stat**). Poor (left): Motor-Right Hand; Mediocre (middle): Emotion: Faces; Good (right): Language: Math-Story. Experiment included 3-100 subjects for training. The group z-statistic baseline results are considerably worse than the group mean baseline. The 4 models largely resemble each other’s performance when sample sizes increase past 40 subjects.

**Figure 8.**
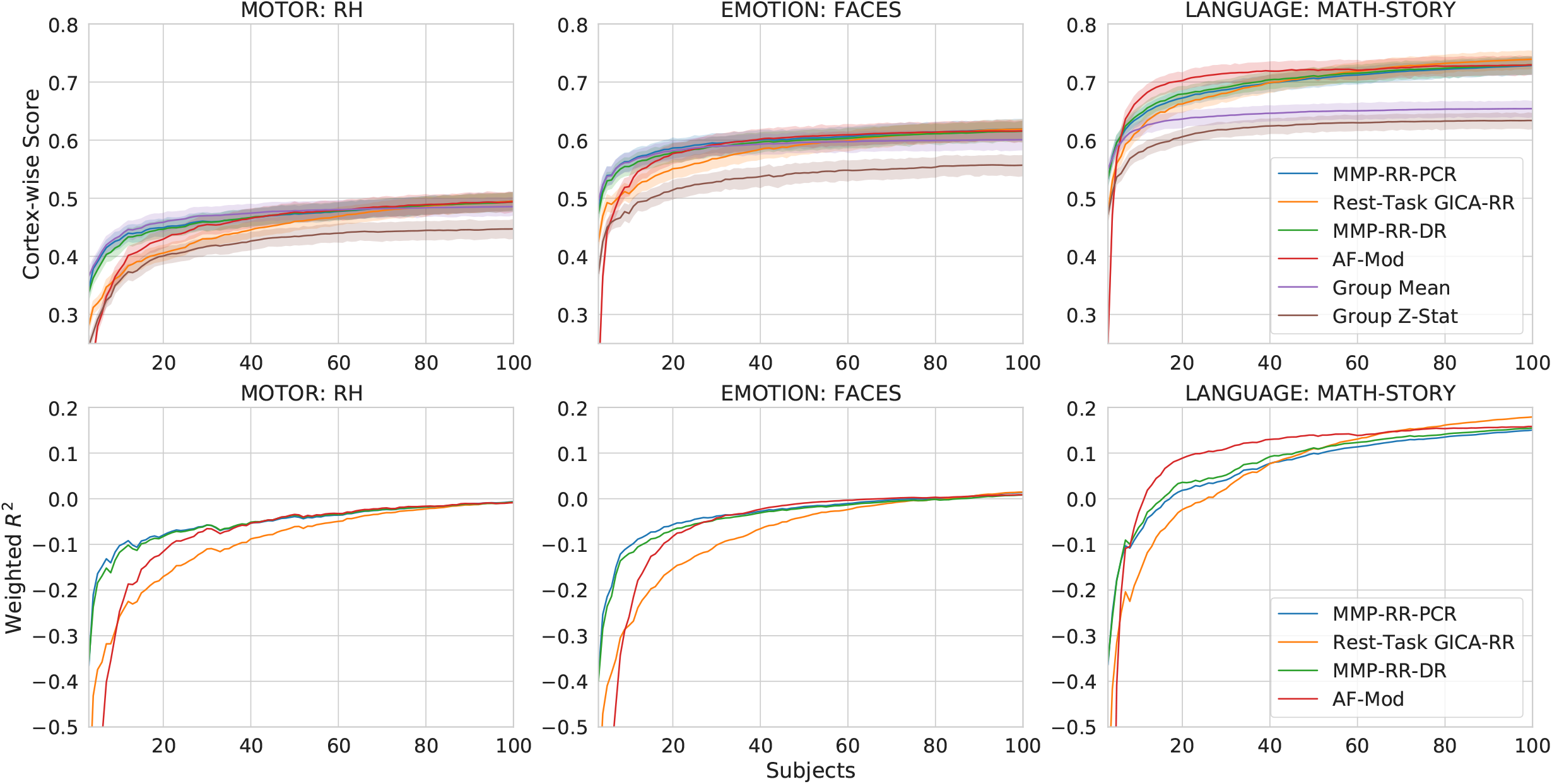
Top row: Subject-wise Pearson *r* score benchmark results for 3 selected (poor, mediocre, good), 4 high performing models (**MMP-RR-PCR**, **Rest-Task GICA-RR**, **MMP-RR-DR**, **AF-Mod**) and two baseline models (**Mean (Baseline)**, **Group Z-stat**). Poor (left): Motor-Right Hand; Mediocre (middle): Emotion: Faces; Good (right): Language: Math-Story. Experiment included 3-100 subjects for training. The group z-statistic baseline results are considerably worse than the group mean baseline. The 4 models largely resemble each other’s performance when sample sizes increase past 40 subjects. Bottom row: The weighted mean cortical surface *R*^2^ for four top performing models as a function of number of samples (3-100) used for training. As training samples approach over 80 samples, *R*^2^ largely becomes indiscernible between the 4 models in these contrasts.

All curves of model performance with respect to the number of samples follow typical generalization curves, i.e., an inverse power law, where a rapid increase is seen to a slow saturation when sample size increases^48^. As shown in both figure 7 and figure 2, **Group Z-stats** consistently underperforms its **Group Mean** counterpart by a considerable margin, especially at lower sample sizes. Top performing models largely yield the same performance as the training set increases above 40 subjects.

### 3.4 Behavioral Results

Prediction scores may provide a powerful means of summarizing rest-task dependency. We therefore hypothesized that prediction scores may be a means for discriminating behaviorally relevant information about the task performed. It was previously speculated that the degree to which brain activity departs from rest may provide information about individual behavioral performance^24^. Within the network neuroscience community, this phenomenon is recognized as reconfiguration efficiency: high-performing individuals may have brain connectivity that more efficiently updates to the task at hand by not having to produce greater changes in a task functional network organization required to perform the task.

We therefore speculated that if our resting-task model performance for individual subjects could be taken as a relative measure of rest-task dependence, we would see a clear pattern of higher behavioral performance correlating with higher tfMRI prediction scores.

To test this idea, we turn to three behavioral measures of general cognitive ability from Human Connectome Data measured during tfMRI acquisition: working memory, language, and reasoning task. We selected contrasts 2BK-0BK, Math-Story, and Match-Relation since they provided the most general and complete summary of the task and its behavioral data. To see whether prediction scores corresponded to task performance of individuals, on the 100-subject test set we calculate the correlation of individual subject Fischer-Z transformed Pearson *r* correlation prediction scores to subject task accuracy. This marks whether individual differences in prediction scores correspond to individual differences in behavioral task accuracy. 20 random train/test permutations of 100 train, 100 test subject sizes on the original 200 subject dataset were fit across the models investigated in the subject-wise investigation. Additionally, similar to figure 7, we also fit the model **MMP-RR-PCR** from these results for the three selected contrasts under 20 permutations train/test splits with increasing sample sizes (3-100 subjects) and expected that these averaged performance evaluation curves would follow typical generalization curves. To accommodate that the 20 training and testing permutations were not independent from each other, statistical comparisons between models were made using a corrected resampled t-test^49^.

Our results demonstrate that Pearson *r* correlation prediction scores provide an indicative relative measure of rest-task correspondence to the behavioral task accuracies measured during the performance of these tasks, figure 9). All predictive models provide statistically significant results over the baseline for contrast Math-Story (one-sided corrected resampled t-test, Fisher-z transformed r, dof=19, *p* < 0.01). Mean correlations over 20 train/test permutations for model **MMP-RR-PCR** compared to **Group Mean** was *r* = 0.26 ± 0.05 versus *r* = 0.20 ± 0.05, respectively. Models **MMP-RR-PCR**, **Rest-Task GICA-RR**, **MMP-RR** provided statistically significant results over baseline for contrast 2BK-0BK (one-sided corrected resampled t-test, Fisher-z tranformed r, dof=19, *p* < 0.05). The mean correlation for contrast 2BK-0BK over 20 train/test permutations for model **MMP-RR-PCR** was *r* = 0.67 ± 0.07 versus **Group Mean** at *r* = 0.66 ± 0.07. However, importantly, in one out of the three contrasts (Match-Relation), no predictive model provides any added benefit over a simple correlation to mean activation (**Group Mean**). That is, despite having strong correlations of *r* = 0.40 ± 0.1. A plot of individual scores for one permutation (original subject test set) is shown in supplementary figure S11 as an illustration of these strong, statistically significant correlations.

**Figure 9.**
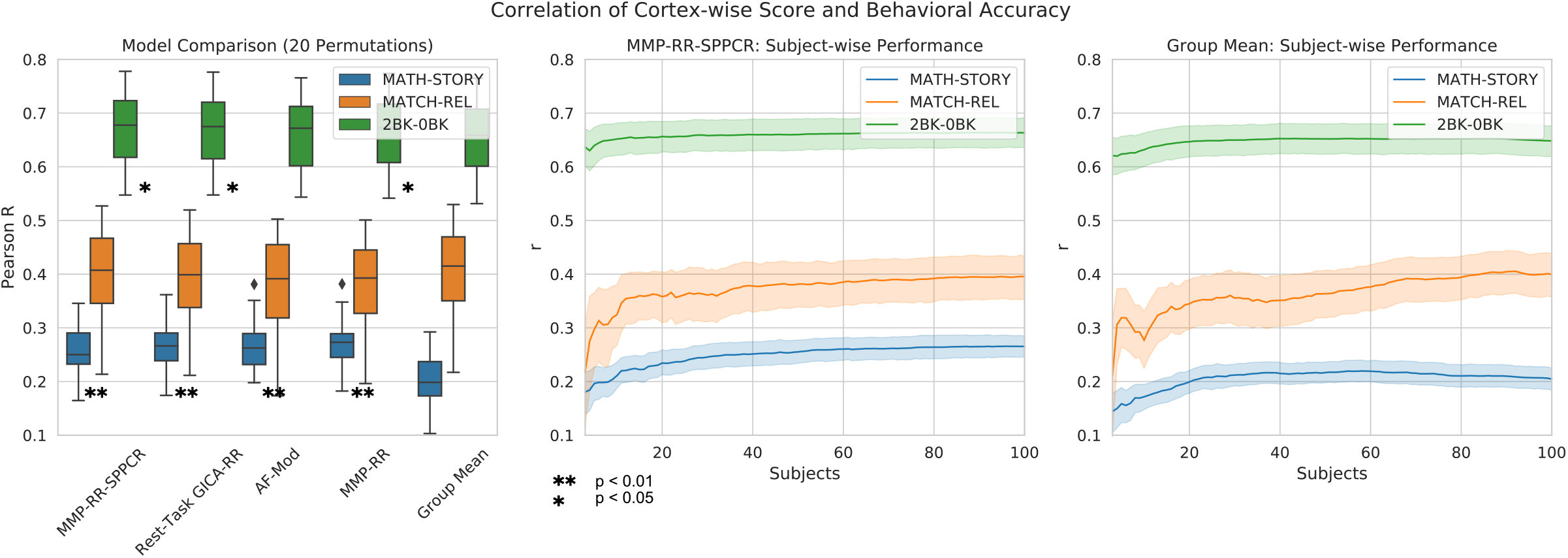
Correlation of Cortex-wise Score and Behavioral Task Accuracy. (A) Comparison of different models investigated in detail shown in figure 8. Only Math-Story and 2BK-0BK showed significant correlations using a one-sided corrected resampled t-test compared to **Group Mean** (significance marked in by *). (B) Subject-wise comparison of correlation between model **MMP-RR-PCR** prediction score and behavioral task performance for 20 permutation depending on the training set size (3-100). (C) For comparison, same as B, but for the **Group Mean** baseline model.

## 4 Discussion

Motivated by recent progress in establishing a stronger link between spontaneous and task-evoked activity, we examine the problem of mapping rsfMRI measurements to patterns of activity elicited during tfMRI-based experimental paradigms in individual subjects. We show additional evidence that it is indeed possible to predict task activity maps from patterns of rsfMRI FC, as previously reported^11–18,23^. However, we emphasized early on that observing higher intra-subject prediction scores compared to inter-subject scores was not a useful observation we believed provided informative predictions–they needed perform better than what any naive group averaging could predict on the cortical surface. Our investigation showed that group averaging provided a surprisingly strong baseline for whole-cortex predictions. Results justify selecting group averaging offered by **Group Mean** as a suitable baseline model. This was because it provides substantially higher scores than its alternative **Group Z-Stat**; group Z-statistics were shown to consistently, regardless of sample size, perform below **Group Mean** under nearly all contrast targets. We therefore evaluated all results against the highest performing baseline–**Group Mean**.

Given this appropriate group-averaged baseline model, an examination of previous methods in our benchmark show they did not demonstrate satisfactory whole-cortex prediction scores with a considerable number of contrast targets being outperformed by the baseline. To remedy this problem, i.e. to jump over the baseline, we introduced a simple modification to the fitting procedure: a vertex-wise selection of hyper-parameters. According to our benchmarks, models fit in this manner provide the most powerful means to tackle the problem of predicting tfMRI GLM maps from rsfMRI data we are aware of. Nevertheless, they also still highlight that in many cases, given the diversity of contrast targets examined, the best performing model we introduced are still modest in their prediction ability with even 100 training samples (subjects).

The considerable variability in prediction scores visible across the 47 contrast maps for all models motivated us to give a better empirical characterization of how this is reflected in model performance. An inspection of the cortical surface areas that have an explainable variance on a vertex-wise level reveals a consistent pattern: primary-sensory regions show little explainable inter-subject variance (*R*^2^) whereas association cortical regions show considerably better predictability. So far, no method appears to be able to explain inter-subject variance within primary sensory regions, as evidenced by strong negative predictive *R*^2^ scores in those locations, figures 5, 6. Additionally, we also observe these patterns by investigating how the strength of regularization was inversely related to how well the model performs. Both measures, *λ* and *R*^2^ shown in figure 6, reveal where information about rsfMRI is actually predictive for task activity maps. Together, these observations reinforce earlier work noting association cortex areas hold distributed networks while primary-sensorimotor areas are much more stereotypical across subjects resulting in worse predictions. Simply put, the closer elicited activity are to these regions–the most salient example being the MATH-STORY contrast–the better these predictions are. Although, many predictions may be better by a statistically significant margin above a baseline model like figure 9 highlight, their utility may still be limited.

Ultimately, our work aims to find which predictions are informative so we could use it to formulate hypotheses asking what behavior or cognitive factors may influence it. That is, the correspondence between rest and task states and how that might reveal information about individual subjects. Seeking to ground this work into a behaviorally relevant context, we considered the question of whether prediction scores of individual subjects provided a means of summarizing rest-task dependence that could inform behaviorally relevant neuroscientific questions given our best performing models. Indeed, the strength of correlation between prediction scores of a given contrast and its corresponding behavioral task accuracy suggests that this prediction score may be taken as a relative measure of dependence between rest and task activity. However, this is not without caveat that places us back to comparing against Group averaged models from the beginning; it is only the case when predictions are considerably above baseline performance we see the utility of performing these model fits. Considerable correlation between the naive model’s prediction of **Group Mean** and individual behavioral performance was present for 2 of the 3 contrasts we examined in this way. This fact reinforces our motivation from the outset of this problem: to create and utilize a method to perform above naive, baseline models. Results shown in our behavior evaluations reiterate this importance. Ultimately, the MATH-STORY fit provided the only meaningful difference compared to the other contrasts examined.

Our vertex-wise evaluation based on predictive *R*^2^ reveals that considerable performance improvement is still needed to explain variance within primary sensorimotor regions. On speculating how to further improve the methods, we suspect that further significant gains in performance may be obtained from projecting individual FC data into common/shared response spaces via shared response modeling or hyperalignment^11,50–53^ This could provide a means for capturing a substantial amount inter-subject variance. Additionally, separate evaluations reveal that the closer the extracted features are to task-related activity, the better cortex-wise prediction scores are, figure S10. We would therefore expect that the use of naturalistic stimuli over rsfMRI could substantially aid over the use of rsfMRI data and would additionally provide the means for additional shared response modeling approach assumptions^54^.

### 4.1 Limitation

First and foremost, should rsfMRI fluctuation amplitudes depend on other factors completely unrelated to cortical computations that generate the spatial dependencies we observe with connectome fingerprinting, this would show up in these prediction result. It would additionally confuse interpretation of behavior factors^55^. Even after application of spatial normalization transformations, considerable anatomical inter-subject variability is preserved despite liberal smoothing application. Additionally, echoplanar imaging (EPI) distortions due to B0 inhomogeneities and other individual specific factors, e.g., coil loading or other RF scaling issues, physiological, motion contaminants, and dependence of individual vascular factors to cortical orientation to B0 would reveal intra-subject dependencies between a rsfMRI and tfMRI acquisition. Regarding the dependence of individual vasculature, large signal biases on BOLD amplitude due to cortical orientations was shown to exist for 3T HCP data^56^. This observation would undoubtedly create additional intra-subject dependencies between measurements that remain after normalization irrespective of any functional organization structure due to underlying neurophysiology or patterns of cortical computations. Therefore, a large degree of dependence will remain after applying normalization transformations and will not necessarily imply that intra-subject prediction scores are necessarily meaningful alone. Disentangling those factors remains to be explored in detail for future work.

Second, the overall test-retest reliability of tfMRI is poor making individual difference research for fMRI difficult with most common task paradigms, especially considering the limited number of task trials GLMs were computed over for HCP data^57^. We would therefore like to emphasize that considerable noise is present in estimates of first-level task effects we sought to predict. In this examination, no model considerations of it was incorporated into any design or analysis decisions.

Last, activity summarized by a task GLM model is a useful measure only insofar as our a priori beliefs about how the task should be parameterized. Encoding models of the task that do not rely on strong assumptions of BOLD response may provide more powerful ways to summarize the kind of dependence we wished to characterize and remains an exciting avenue to explore beyond GLM maps^58^.

## 5 Summary

Our closer examination using Human Connectome Project (HCP) data reveals that a majority of published models evaluated within our benchmark under current methods with many contrast targets examined did not perform better than naive, baseline models when only rsfMRI features and whole-cortex prediction were considered. This paper aims to remedy this issue and make a convincing case for utilizing methods to describe individual factors beyond merely remarking on individual differences. We propose single-vertex fitted methods that achieve a significant performance boost above baseline performance on the majority of contrast targets. Additionally, we provide benchmarks of comparable methods in published literature and include a variety of models with feature properties worth investigating, table 1. We provide further empirical characterization of top performing methods by an examination of showing where predictions performed well spatially. This explains why predictions of only a modest number of contrasts is possible above a naive baseline. Ultimately, we show that a model’s prediction score can be taken as a relative measure of dependency between rest and task. These predictions results show a compelling behavioral correspondence to a subject’s task performance committed during a tfMRI acquisition albeit with notable caveats. We hope that further improvements to this methodology will enable better understanding of rest-task correspondence informing individual behavioral measures.

## 6 Acknowledgements

Data were provided by the Human Connectome Project, WU-Minn Consortium (Principal Investigators: David Van Essen and Kamil Ugurbil; 1U54MH091657) funded by the 16 NIH Institutes and Centers that support the NIH Blueprint for Neuroscience Research; and by the McDonnell Center for Systems Neuroscience at Washington University.

