## Supplementary for "Jumping over Baselines with New Methods to Predict Activation Maps from Resting-state fMRI"

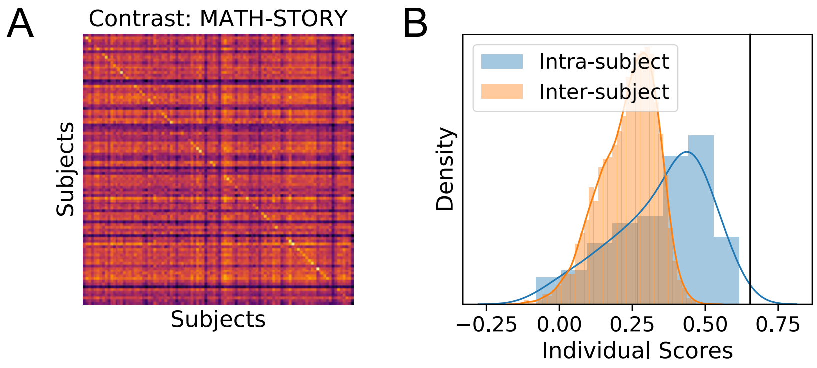


**Suppl. Figure S1.** Higher intra-subject than inter-subject scores for task activation map prediction can be shown with minimal modeling and requiring no statistical fitting. We illustrate this with an arbitrary example under the the Math-Story contrast target. Here, a random projection (see rsfMRI feature extraction section) where $A$ is sampled from $\mathcal{N}(0,1)$ can demonstrate a higher intra-subject than inter-subject scores to a subject's task activation map. (A) shows the subject-wise confusion matrix. (B) shows a density plot of intra-subject vs. inter-subject scores. Brains measured from the same subject with rsfMRI and tfMRI are more similar to each other than different subjects. These scores are still considerably worse than a **Group Mean** baseline marked in the solid black line.

### Supplementary Information

#### Model Evaluation

Dice overlaps and higher intra-subject than inter-subject scores have been widely used in related literature to present support for successful model predictions performance. We believe reliance on these metrics can be highly misleading and have therefore avoided its use. This is illustrated with two examples that highlight this. First, figure S1 shows how higher intra-subject than inter-subject scores can be trivially obtained to show individual differences. Second, Dice coefficients have properties that make the metric poor for evaluating model performance. This is based on two factors: (1) sensitivity to chosen thresholds that change baseline and fitted models performance differently shown in S2 and (2) non-intuitive biases that occur as a result of increasing the number of subjects used calculation of baseline models, lowering model performance, especially at conservative thresholds shown in S3.

##### Group Z-Stat (TFCE) details

Calculation of **Group Z-Stat (TFCE)** was performed as follows: group-level statistical inference was calculated within CIFTI-greyordinate space form via Permutation Analysis of Linear Models (PALM)[1] that incorporates spatial statistics for Threshold Free Cluster Enhancement (TFCE)[2]. This was done by performing inference separately for each brain hemisphere and CIFTI volumetric component since surface and volume space yield different spatial statistics used by TFCE. Surface meshes to incorporate these spatial statistics were computed as the mean mid-thickness cortical surface across all 200 subjects via the Connectome Workbench toolbox[3] `wb_command -surface-vertex-areas` and then merged together `wb_command -metric-merge` and finally averaged `wb_command -metric-reduce`. Group-level one-sample t-tests were computed ignoring intra-subject variance of contrast parameter estimates for all 47 contrasts examined. To facilitate quicker computation times, p-value estimates were computed via a Gamma fit approximation with 500 permutations performed for the null-distribution estimate of each contrast assuming independent and symmetric errors as implemented in PALM[1]. Group-level maps were finally transformed to a z-statistic and thresholded at $p < 0.05$ (FWER-corrected for multiple comparisons using TFCE).

#### Additional Subject-wise Evaluation

To discriminate more closely between models in the subject-wise evaluation, the plot figure 8 offers a clearer view offered by the $R^2$ evaluation metric. However, in order to better differentiate between them relative to their sample sizes, the fraction of cortical surfaces having an $R^2$ above some given threshold is plotted. 3 different thresholds are included to provide the greatest discriminability between models at thresholds $0, 0.1, 0.2$. The sensitivity of the vertex-wise $R^2$ score from the chosen threshold is plotted in supplementary figure S8. For the 3 contrasts examined, at lowest thresholds, model **AF-mod** shows greatest fractional $R^2$ above 0 for all 3 contrasts while **MMP-RR-PCR** shows greatest fraction $R^2$ for at least 2/3 contrasts,

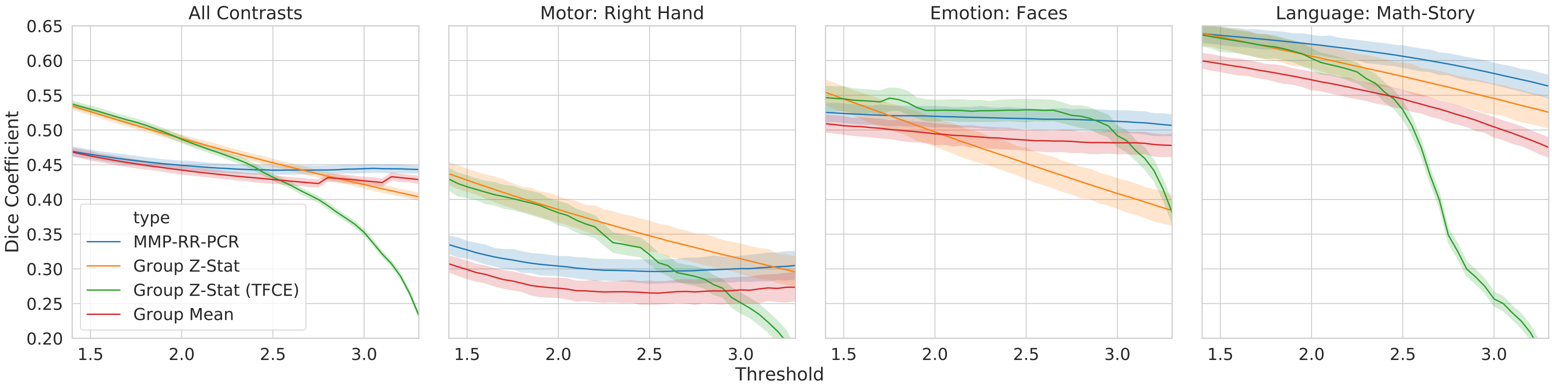


**Suppl. Figure S2.** Dice score sensitivity to choosen threshold: Single subject GLM maps hold a considerable degree of noise resulting in inflated statistical errors in detecting activation in task-based fMRI. Therefore, many results reported on the vertex/voxel level are often done on a binary active vs. non-active classification basis and are sometimes utilized to evaluate fMRI reproducibility measures. Here, we report on the stability of applying a threshold procedure for the model evaluation. Varying thresholds (Z-scores: 1.4-3.3) were chosen to display the overall sensitivity to results over baseline models: Group Z-stat, Group Z-stat Threshold Free Cluster Extent (TFCE) corrected at Family-wise Error (FWE) $p < 0.05$, and Group Mean. Liberal thresholds show Dice coefficient scores higher for group-based models than a top-performing fitted. Fitted models, according to this evaluation metric, show better performance than **Group** models only when thresholds are increasingly strict.

especially at the highest threshold 0.2. Differences between these models are seen depending on number of training subjects, threshold selected, and contrast examined.

#### Miscellaneous

#### 1 Score Table

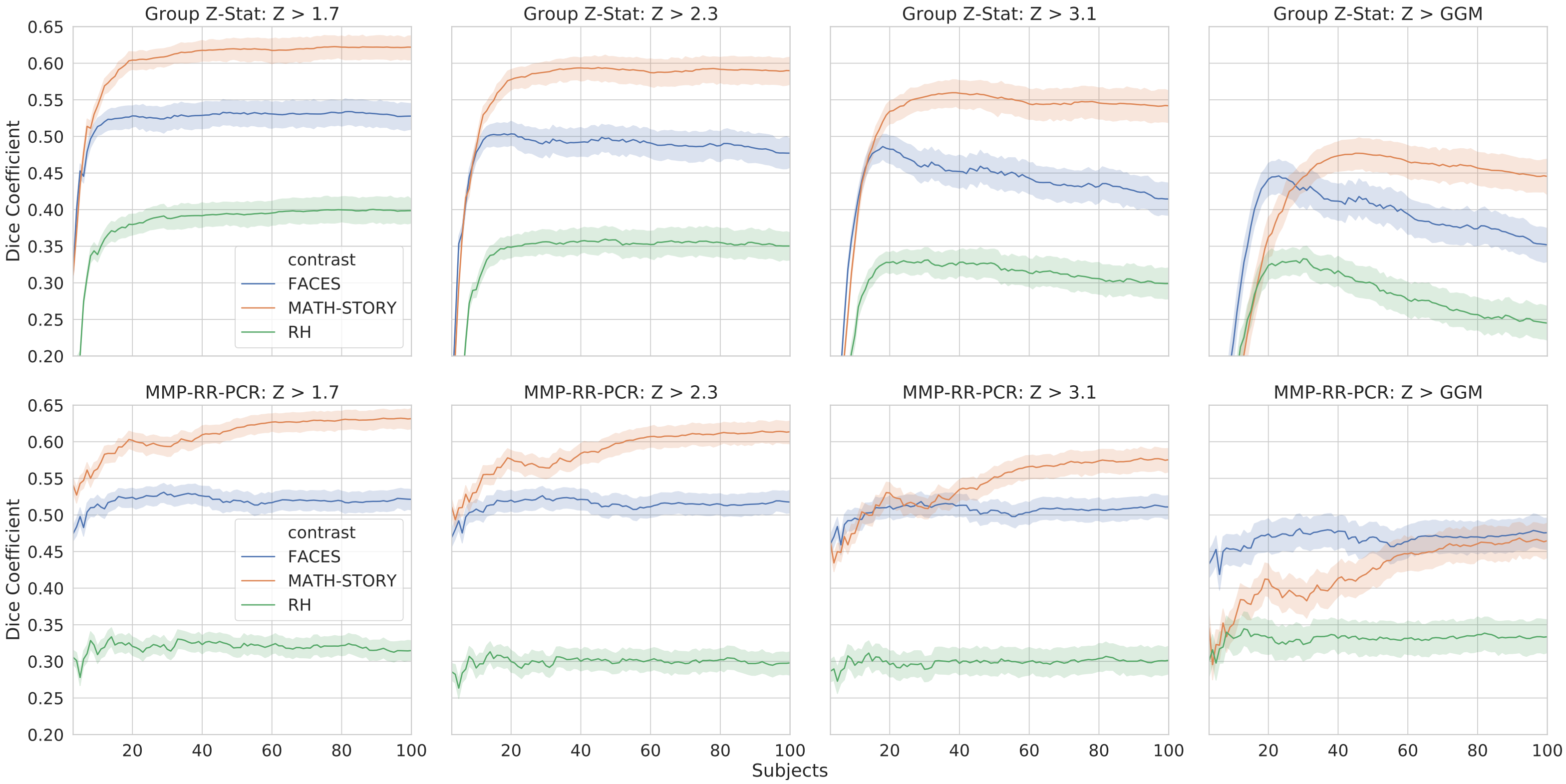


**Suppl. Figure S3.** Dice score sensitivity to number of training subjects: A comparison between **Group Z-Stat** models (top row) vs. **MMP-RR-PCR** (bottom row) with varying samples (subjects) used to fit the models. Shown are 3 different contrasts used in the main text for subject-wise evaluation. Column plots show increasing (left to right) Z thresholds choosen for the comparison, i.e., 1.7, 2.3, 3.1, and GGM–a threshold calculated from individually fitted Gaussian Gamma Mixture models on actual task activation maps of individual subjects, which tend to be considerably more conservative than $Z > 3.1$. This threshold was the estimated median of the positive Gamma component. Higher thresholds show that an increasing number of subjects used to calculate **Group Z-Stat** models lowers Dice score considerably. For fit model MMP-RR-PCR on contrasts FACES or RH, the fitted contrasts appear to remain flat across all thresholds without considerable and expected increases in Dice score due to increasing sample size.

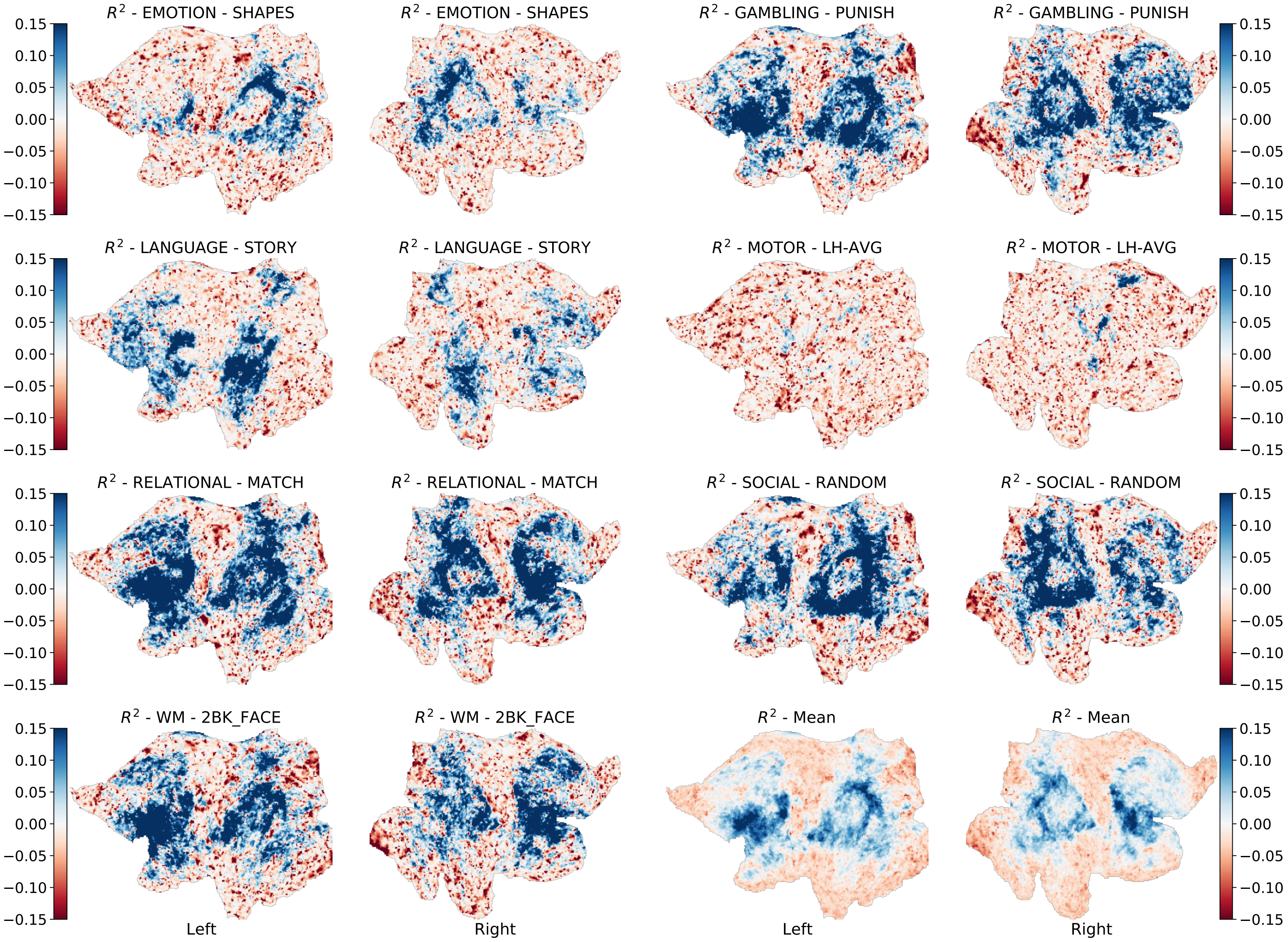


**Suppl. Figure S4.** Separate test-sample $R^2$ of model **MMP-RR-PCR** evaluations of a single task contrasts belonging to each task category: All 47 contrast targets belong to 7 different task categories: Emotion, Gambling, Language, Motor, Relational, Social, Working Memory. A contrast in each category closest to the median Pearson R score of that category was selected for display. The test-sample mean over all 47 contrasts is plotted also for convenience. A general pattern is clear across all contrasts: it is only within certain regions, e.g., association cortex, that a positive $R^2$ appears possible. Primary sensorimotor regions are consistently negative.

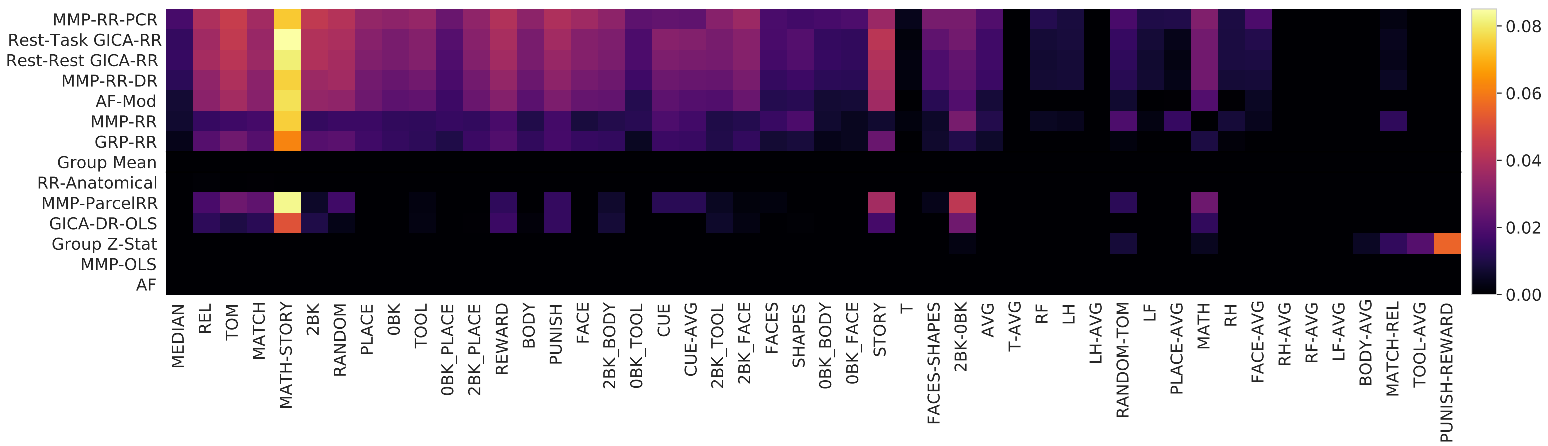


**Suppl. Figure S5.** Pearson $r$ correlation score benchmark results for 100 subject test set: Colorbar indicates mean $r$ score difference between the model prediction score and the mean baseline across all test subjects for given contrast and model. Scores are ordered by model and contrast exactly like figure 2. This figure is akin to figure 4 showing what models achieve an $R^2$ score above 0.

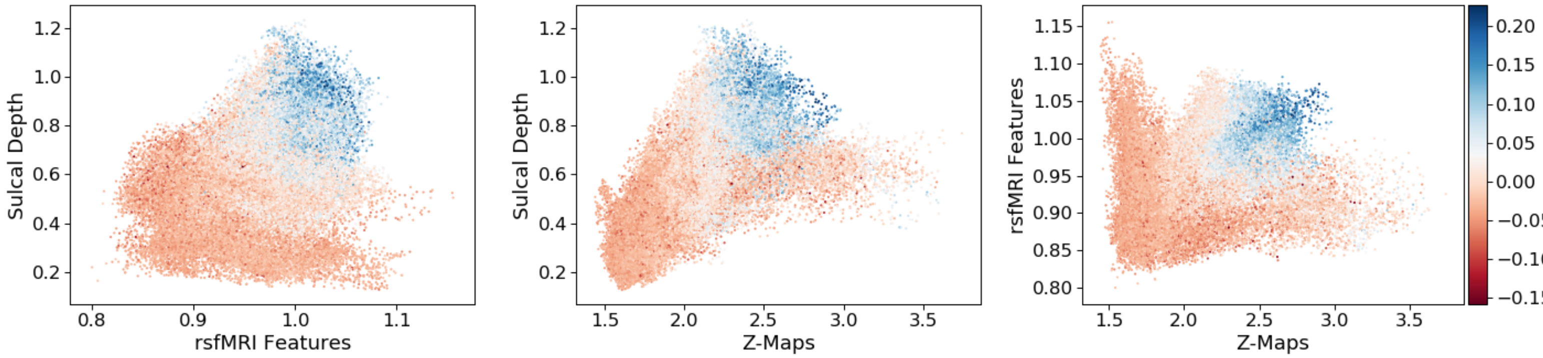


**Suppl. Figure S6.** Inter-subject variability of activation maps (Z-Maps), sulcal depth, or **MMP-RR-PCR** rsfMRI features and how they relate to measured $R^2$ and each other plotted as a 3D scatter plot where each point represents a cortical vertex. The colormap represents a vertices' mean $R^2$ computed across all 47 contrasts. Inter-subject variability of features, sulcal depth, or Z-maps are computed as the vertex-wise standard deviations. rsfMRI features and Z-Maps are averaged across all features (379) or contrasts (47). This plot shows sulcul depth and rsfMRI functional correlational features are strongly correlated to one another. Unsurprisingly, the model's prediction ability as measured by $R^2$ on a vertex-wise basis are concentrated around the point-cloud mass where inter-subject variability between the two factors are highest (upper right-most areas of the plots).

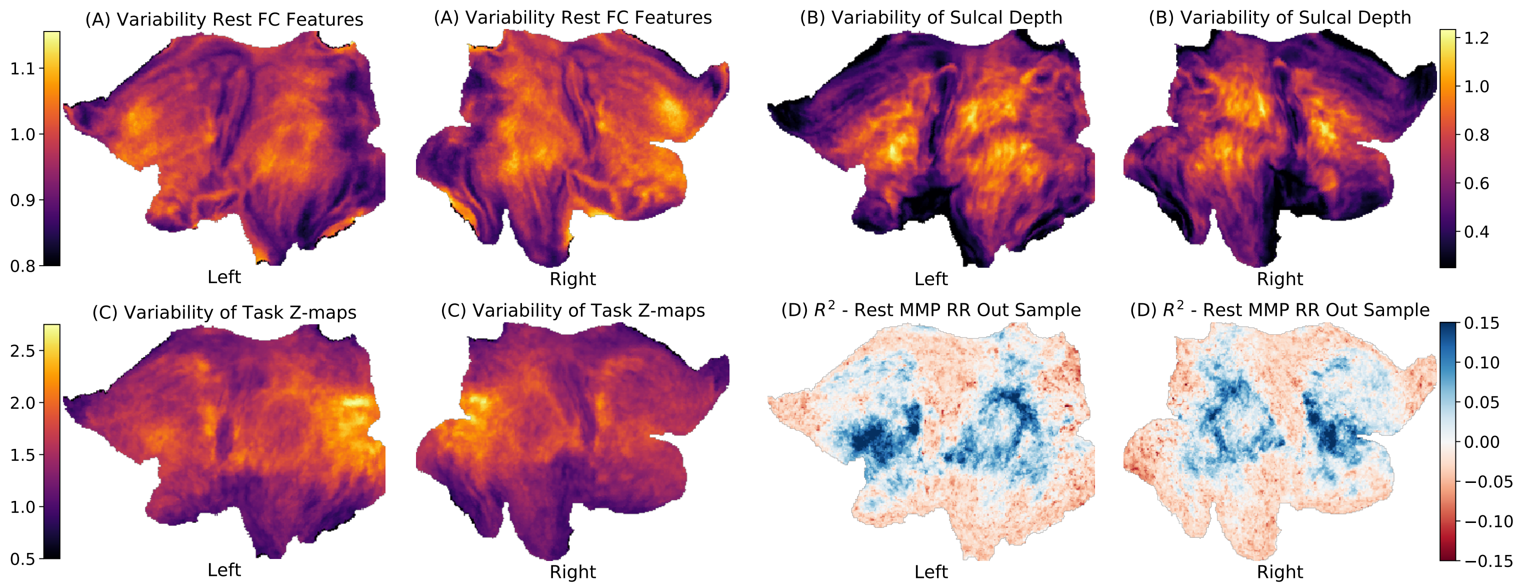


**Suppl. Figure S7.** Flatmap projections of vertex-wise inter-subject variability as shown in figure S6. Shown is (A) **MMP-RR-PCR** FC Features, (B) sulcal depth, (C) task activation Z-maps. (D) shows predictive $R^2$ of model **MMP-RR-PCR**.

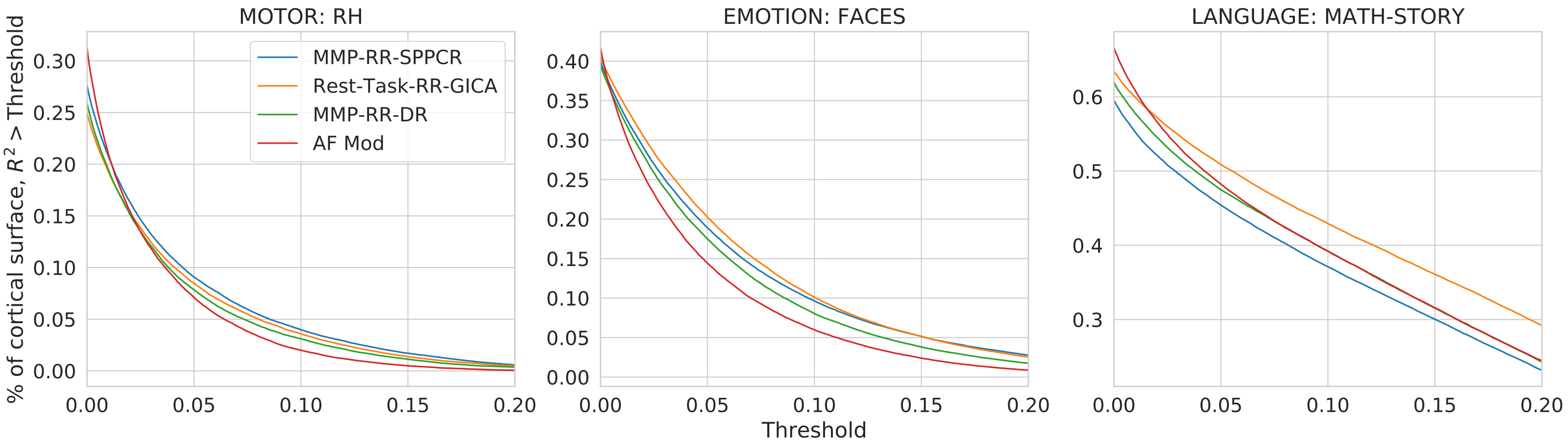


**Suppl. Figure S8.** Vertex-wise $R^2$ score sensitivity at 100 training samples: scores indicate the fraction of cortical surface having an $R^2$ score above a given threshold plotted on the x-axis for given contrast and model.

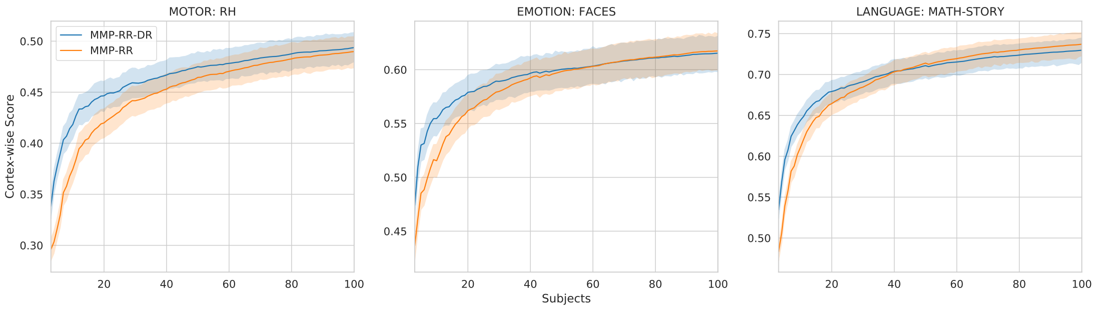


**Suppl. Figure S9.** **MMP-RR-DR** and **MMP-RR** subject-wise comparison. Applying Dual Regression for feature extraction yielded better vertex-wise and cortex-wise scores for a smaller number of training samples, around less than 40. However, the difference between the two models were negligible when training samples increased to 100. We speculate that dual regression might offer the most benefits with small subject sizes.

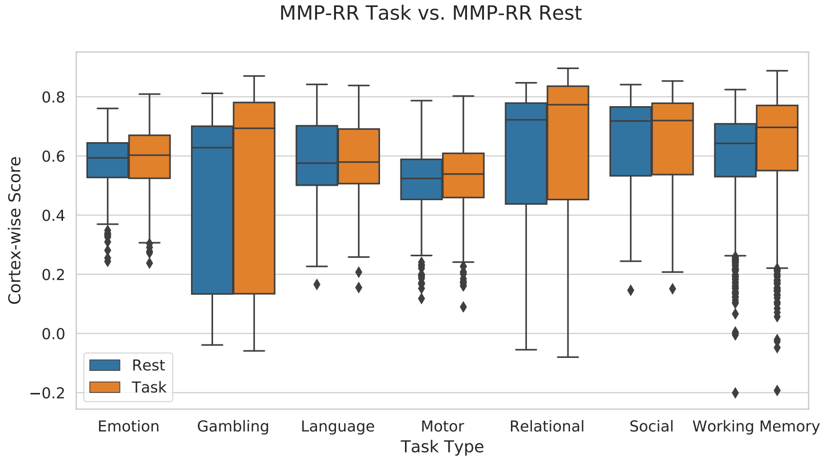


**Suppl. Figure S10.** We investigated whether task data from separate task measurements provided better features for task prediction than resting-state data. Implicitly it is assumed resting-state scans, as opposed to task-based data where a very limited number of cognitive brain activity modes are investigated, subjects will enter multiple cognitive modes comprising of default, visual, motor, executive control and attention processes. This is supported by the networks of brain activity that are elicited during a single measurement of rest largely overlapping those that are extracted during the task[4,5]. Furthermore, even across different task states, it appears that a core cognitive network dominates[6]. Do these multiple cognitive modes that are speculatively elicited during a rest scan differentiate subjects better than tfMRI?. Here, we tested whether a concatenation of data from the HCP battery of 6 diverse, but ultimately limited tasks[7]. In the tfMRI case, separate features were calculated by selecting only 6 of the 7 tfMRI datasets, leaving out the tfMRI measurement of the to-be predicted GLM task contrast. Doing this excluded circularity. In reported experiments, data matrix $X_i$ was a concatenation across 6 of the 7 task measurements with the excluded measurement being the one under which the contrast map was computed with. This led to features being computed from 3468, 3314, 3188, 3252, 3356, 3272, 3010 samples for EMOTION, GAMBLING, LANGUAGE, MOTOR, RELATIONAL, SOCIAL, WM contrast map predictions, respectively.)

| Contrast | MMP-RR-PCR | Rest-Task GICA-RR | Rest-Rest GICA-RR | MMP-RR-DR | MMP-RR | AF-Mod | GRP-RR | Group Mean | RR-Anatomical | MMP-ParcelRR | GICA-DR-OLS | Group Z-Stat | MMP-OLS | AF |
|---|---|---|---|---|---|---|---|---|---|---|---|---|---|---|
| MEAN | 0.582 | 0.581 | 0.58 | 0.578 | 0.574 | 0.571 | 0.568 | 0.561 | 0.557 | 0.551 | 0.55 | 0.54 | 0.409 | 0.302 |
| REL | 0.766 | 0.763 | 0.764 | 0.759 | 0.759 | 0.741 | 0.748 | 0.726 | 0.727 | 0.745 | 0.74 | 0.693 | 0.639 | 0.46 |
| TOM | 0.765 | 0.764 | 0.763 | 0.76 | 0.758 | 0.737 | 0.747 | 0.721 | 0.72 | 0.747 | 0.731 | 0.685 | 0.634 | 0.421 |
| MATCH | 0.751 | 0.749 | 0.749 | 0.745 | 0.745 | 0.731 | 0.734 | 0.713 | 0.714 | 0.736 | 0.726 | 0.679 | 0.621 | 0.479 |
| MATH-STORY | 0.729 | 0.739 | 0.735 | 0.73 | 0.732 | 0.729 | 0.716 | 0.654 | 0.654 | 0.737 | 0.705 | 0.642 | 0.609 | 0.618 |
| 2BK | 0.739 | 0.735 | 0.735 | 0.73 | 0.729 | 0.709 | 0.716 | 0.695 | 0.694 | 0.701 | 0.705 | 0.666 | 0.604 | 0.397 |
| RANDOM | 0.733 | 0.731 | 0.73 | 0.729 | 0.725 | 0.708 | 0.714 | 0.692 | 0.69 | 0.708 | 0.695 | 0.654 | 0.585 | 0.433 |
| PLACE | 0.735 | 0.732 | 0.731 | 0.728 | 0.727 | 0.717 | 0.718 | 0.701 | 0.7 | 0.692 | 0.698 | 0.65 | 0.594 | 0.4 |
| 0BK | 0.723 | 0.718 | 0.718 | 0.715 | 0.713 | 0.704 | 0.704 | 0.69 | 0.689 | 0.677 | 0.687 | 0.64 | 0.577 | 0.375 |
| TOOL | 0.712 | 0.708 | 0.708 | 0.704 | 0.701 | 0.691 | 0.691 | 0.678 | 0.676 | 0.68 | 0.681 | 0.644 | 0.562 | 0.399 |
| 0BK_PLACE | 0.692 | 0.688 | 0.687 | 0.686 | 0.684 | 0.682 | 0.678 | 0.667 | 0.666 | 0.651 | 0.658 | 0.624 | 0.53 | 0.391 |
| 2BK_PLACE | 0.693 | 0.691 | 0.69 | 0.687 | 0.685 | 0.674 | 0.676 | 0.66 | 0.658 | 0.658 | 0.661 | 0.628 | 0.539 | 0.368 |
| REWARD | 0.687 | 0.685 | 0.684 | 0.681 | 0.677 | 0.665 | 0.667 | 0.647 | 0.645 | 0.66 | 0.662 | 0.625 | 0.544 | 0.412 |
| BODY | 0.683 | 0.679 | 0.679 | 0.676 | 0.672 | 0.661 | 0.664 | 0.65 | 0.649 | 0.649 | 0.652 | 0.627 | 0.525 | 0.358 |
| PUNISH | 0.669 | 0.666 | 0.663 | 0.662 | 0.658 | 0.646 | 0.647 | 0.629 | 0.627 | 0.643 | 0.643 | 0.613 | 0.517 | 0.395 |
| FACE | 0.676 | 0.67 | 0.67 | 0.667 | 0.664 | 0.649 | 0.654 | 0.639 | 0.638 | 0.624 | 0.633 | 0.608 | 0.518 | 0.304 |
| 2BK_BODY | 0.662 | 0.658 | 0.658 | 0.655 | 0.652 | 0.64 | 0.643 | 0.629 | 0.627 | 0.636 | 0.637 | 0.618 | 0.502 | 0.363 |
| 0BK_TOOL | 0.653 | 0.65 | 0.65 | 0.647 | 0.642 | 0.643 | 0.636 | 0.631 | 0.628 | 0.627 | 0.627 | 0.602 | 0.476 | 0.367 |
| CUE | 0.643 | 0.651 | 0.649 | 0.645 | 0.642 | 0.639 | 0.635 | 0.62 | 0.619 | 0.632 | 0.612 | 0.572 | 0.478 | 0.421 |
| CUE-AVG | 0.644 | 0.651 | 0.649 | 0.646 | 0.642 | 0.638 | 0.636 | 0.621 | 0.621 | 0.633 | 0.617 | 0.552 | 0.489 | 0.405 |
| 2BK_TOOL | 0.65 | 0.647 | 0.646 | 0.643 | 0.639 | 0.629 | 0.629 | 0.619 | 0.616 | 0.624 | 0.625 | 0.604 | 0.482 | 0.362 |
| 2BK_FACE | 0.654 | 0.65 | 0.649 | 0.647 | 0.643 | 0.63 | 0.632 | 0.618 | 0.616 | 0.62 | 0.621 | 0.607 | 0.49 | 0.338 |
| FACES | 0.619 | 0.619 | 0.618 | 0.615 | 0.612 | 0.616 | 0.609 | 0.601 | 0.597 | 0.603 | 0.592 | 0.57 | 0.435 | 0.397 |
| SHAPES | 0.597 | 0.601 | 0.6 | 0.596 | 0.592 | 0.599 | 0.589 | 0.58 | 0.576 | 0.561 | 0.581 | 0.559 | 0.405 | 0.417 |
| 0BK_BODY | 0.586 | 0.582 | 0.583 | 0.58 | 0.576 | 0.575 | 0.572 | 0.568 | 0.564 | 0.558 | 0.561 | 0.552 | 0.396 | 0.304 |
| 0BK_FACE | 0.575 | 0.57 | 0.57 | 0.568 | 0.564 | 0.56 | 0.56 | 0.556 | 0.553 | 0.522 | 0.537 | 0.528 | 0.386 | 0.233 |
| STORY | 0.549 | 0.556 | 0.554 | 0.552 | 0.551 | 0.521 | 0.539 | 0.514 | 0.511 | 0.551 | 0.532 | 0.511 | 0.38 | 0.186 |
| T | 0.534 | 0.531 | 0.532 | 0.532 | 0.526 | 0.532 | 0.527 | 0.529 | 0.526 | 0.484 | 0.502 | 0.513 | 0.327 | 0.243 |
| FACES-SHAPES | 0.545 | 0.539 | 0.536 | 0.536 | 0.528 | 0.523 | 0.523 | 0.516 | 0.515 | 0.52 | 0.492 | 0.474 | 0.366 | 0.221 |
| 2BK-0BK | 0.512 | 0.512 | 0.508 | 0.507 | 0.505 | 0.513 | 0.495 | 0.484 | 0.479 | 0.528 | 0.511 | 0.487 | 0.327 | 0.402 |
| AVG | 0.53 | 0.527 | 0.527 | 0.526 | 0.519 | 0.521 | 0.517 | 0.51 | 0.508 | 0.491 | 0.494 | 0.493 | 0.341 | 0.262 |
| T-AVG | 0.531 | 0.529 | 0.53 | 0.53 | 0.523 | 0.533 | 0.525 | 0.533 | 0.528 | 0.454 | 0.497 | 0.498 | 0.309 | 0.197 |
| RF | 0.52 | 0.517 | 0.516 | 0.516 | 0.508 | 0.514 | 0.507 | 0.508 | 0.505 | 0.472 | 0.48 | 0.49 | 0.327 | 0.201 |
| LH | 0.516 | 0.516 | 0.515 | 0.515 | 0.504 | 0.512 | 0.506 | 0.507 | 0.504 | 0.457 | 0.482 | 0.484 | 0.317 | 0.213 |
| LH-AVG | 0.517 | 0.519 | 0.518 | 0.518 | 0.51 | 0.516 | 0.51 | 0.52 | 0.516 | 0.453 | 0.487 | 0.503 | 0.3 | 0.09 |
| RANDOM-TOM | 0.496 | 0.492 | 0.49 | 0.489 | 0.484 | 0.497 | 0.479 | 0.477 | 0.474 | 0.49 | 0.466 | 0.485 | 0.301 | 0.353 |
| LF | 0.513 | 0.51 | 0.509 | 0.509 | 0.502 | 0.505 | 0.502 | 0.502 | 0.498 | 0.468 | 0.471 | 0.485 | 0.313 | 0.185 |
| PLACE-AVG | 0.504 | 0.497 | 0.496 | 0.496 | 0.493 | 0.507 | 0.489 | 0.493 | 0.487 | 0.471 | 0.443 | 0.489 | 0.296 | 0.281 |
| MATH | 0.504 | 0.501 | 0.501 | 0.501 | 0.495 | 0.473 | 0.484 | 0.474 | 0.471 | 0.501 | 0.488 | 0.479 | 0.327 | 0.122 |
| RH | 0.495 | 0.495 | 0.495 | 0.494 | 0.486 | 0.494 | 0.487 | 0.486 | 0.482 | 0.444 | 0.463 | 0.465 | 0.294 | 0.206 |
| FACE-AVG | 0.495 | 0.487 | 0.486 | 0.485 | 0.481 | 0.481 | 0.476 | 0.476 | 0.469 | 0.473 | 0.424 | 0.469 | 0.282 | 0.291 |
| RH-AVG | 0.486 | 0.487 | 0.487 | 0.486 | 0.48 | 0.487 | 0.481 | 0.492 | 0.486 | 0.419 | 0.458 | 0.469 | 0.267 | 0.115 |
| RF-AVG | 0.466 | 0.467 | 0.466 | 0.465 | 0.46 | 0.468 | 0.462 | 0.474 | 0.468 | 0.403 | 0.415 | 0.448 | 0.248 | 0.092 |
| LF-AVG | 0.465 | 0.463 | 0.462 | 0.463 | 0.457 | 0.466 | 0.458 | 0.472 | 0.466 | 0.4 | 0.413 | 0.435 | 0.245 | 0.055 |
| BODY-AVG | 0.398 | 0.394 | 0.395 | 0.394 | 0.388 | 0.392 | 0.388 | 0.398 | 0.392 | 0.369 | 0.356 | 0.404 | 0.196 | 0.108 |
| MATCH-REL | 0.353 | 0.354 | 0.354 | 0.356 | 0.346 | 0.363 | 0.348 | 0.35 | 0.341 | 0.333 | 0.336 | 0.364 | 0.168 | 0.269 |
| TOOL-AVG | 0.311 | 0.305 | 0.304 | 0.304 | 0.295 | 0.3 | 0.291 | 0.315 | 0.303 | 0.294 | 0.281 | 0.336 | 0.131 | 0.176 |
| PUNISH-REWARD | 0.096 | 0.096 | 0.096 | 0.095 | 0.093 | 0.102 | 0.093 | 0.118 | 0.105 | 0.086 | 0.091 | 0.173 | 0.026 | 0.086 |

**Suppl. Table S1.** Correlation Score Table: Mean Correlation Scores averaged over subjects and shown across all models examined ordered by Contrast names as in figure 2. Additional supplementary material provides individual subject scores in a CSV (all_model_and_subject_r_scores.csv).

| Contrast | MMP-RR-PCR | Rest-Task GICA-RR | Rest-Rest GICA-RR | MMP-RR-DR | MMP-RR | AF-Mod | GRP-RR | Group Mean | RR-Anatomical | MMP-ParcelRR | GICA-DR-OLS | Group Z-Stat | MMP-OLS | AF |
|---|---|---|---|---|---|---|---|---|---|---|---|---|---|---|
| MEAN | 0.025 | 0.022 | 0.02 | 0.018 | 0.011 | 0.001 | -0.002 | -0.02 | -0.098 | -0.029 | -0.076 | -2.649 | -0.686 | -940632.771 |
| REL | 0.095 | 0.086 | 0.088 | 0.08 | 0.078 | 0.022 | 0.04 | -0.021 | -0.087 | 0.03 | -0.062 | -3.204 | -0.537 | -2297521.704 |
| TOM | 0.107 | 0.102 | 0.096 | 0.094 | 0.083 | 0.024 | 0.048 | -0.022 | -0.09 | 0.045 | -0.056 | -3.258 | -0.511 | -4063035.33 |
| MATCH | 0.081 | 0.071 | 0.073 | 0.067 | 0.059 | 0.025 | 0.028 | -0.022 | -0.095 | 0.039 | -0.066 | -3.48 | -0.596 | -2716221.733 |
| MATH-STORY | 0.151 | 0.179 | 0.168 | 0.155 | 0.166 | 0.158 | 0.124 | -0.019 | -0.085 | 0.157 | 0.061 | -1.963 | -0.336 | -282755.818 |
| 2BK | 0.092 | 0.083 | 0.083 | 0.072 | 0.074 | 0.015 | 0.035 | -0.024 | -0.095 | -0.014 | -0.076 | -1.045 | -0.497 | -1182910.045 |
| RANDOM | 0.081 | 0.077 | 0.074 | 0.075 | 0.065 | 0.019 | 0.033 | -0.02 | -0.093 | 0.02 | -0.073 | -2.771 | -0.545 | -4061690.458 |
| PLACE | 0.07 | 0.06 | 0.057 | 0.051 | 0.05 | 0.019 | 0.022 | -0.022 | -0.094 | -0.052 | -0.103 | -1.867 | -0.589 | -965346.968 |
| 0BK | 0.06 | 0.048 | 0.048 | 0.041 | 0.038 | 0.011 | 0.012 | -0.023 | -0.094 | -0.063 | -0.094 | -1.192 | -0.583 | -1250291.134 |
| TOOL | 0.062 | 0.054 | 0.053 | 0.045 | 0.043 | 0.011 | 0.014 | -0.021 | -0.1 | -0.023 | -0.079 | -1.494 | -0.581 | -1047548.118 |
| 0BK_PLACE | 0.038 | 0.026 | 0.024 | 0.022 | 0.018 | 0.012 | 0.002 | -0.021 | -0.093 | -0.063 | -0.094 | -1.975 | -0.664 | -546003.116 |
| 2BK_PLACE | 0.052 | 0.047 | 0.046 | 0.039 | 0.035 | 0.009 | 0.012 | -0.023 | -0.101 | -0.032 | -0.091 | -2.09 | -0.627 | -1033948.366 |
| REWARD | 0.067 | 0.061 | 0.057 | 0.057 | 0.049 | 0.019 | 0.023 | -0.02 | -0.094 | 0.007 | -0.048 | -3.146 | -0.539 | -976646.385 |
| BODY | 0.047 | 0.039 | 0.04 | 0.033 | 0.028 | 0.0 | 0.007 | -0.023 | -0.094 | -0.033 | -0.08 | -1.613 | -0.593 | -1429015.359 |
| PUNISH | 0.06 | 0.055 | 0.048 | 0.049 | 0.042 | 0.014 | 0.016 | -0.021 | -0.098 | 0.006 | -0.056 | -3.31 | -0.588 | -666174.275 |
| FACE | 0.05 | 0.038 | 0.04 | 0.033 | 0.03 | -0.007 | 0.007 | -0.027 | -0.102 | -0.062 | -0.114 | -1.384 | -0.595 | -732123.083 |
| 2BK_BODY | 0.046 | 0.04 | 0.039 | 0.034 | 0.032 | 0.001 | 0.011 | -0.021 | -0.09 | -0.014 | -0.056 | -2.221 | -0.562 | -877546.337 |
| 0BK_TOOL | 0.024 | 0.016 | 0.017 | 0.012 | 0.005 | 0.001 | -0.011 | -0.022 | -0.113 | -0.035 | -0.077 | -1.87 | -0.738 | -825311.656 |
| CUE | 0.027 | 0.044 | 0.041 | 0.035 | 0.02 | 0.019 | 0.002 | -0.017 | -0.089 | 0.014 | -0.098 | -9.776 | -0.688 | -5853287.768 |
| CUE-AVG | 0.025 | 0.038 | 0.035 | 0.031 | 0.011 | 0.012 | -0.005 | -0.019 | -0.096 | 0.013 | -0.097 | -11.297 | -0.682 | -5293763.983 |
| 2BK_TOOL | 0.039 | 0.033 | 0.032 | 0.027 | 0.023 | -0.001 | 0.002 | -0.021 | -0.1 | -0.016 | -0.065 | -1.713 | -0.621 | -890898.033 |
| 2BK_FACE | 0.046 | 0.038 | 0.038 | 0.033 | 0.03 | -0.001 | 0.006 | -0.024 | -0.104 | -0.023 | -0.086 | -1.661 | -0.61 | -480804.67 |
| FACES | 0.014 | 0.015 | 0.012 | 0.008 | 0.001 | 0.009 | -0.008 | -0.019 | -0.101 | -0.013 | -0.069 | -3.65 | -0.746 | -880524.147 |
| SHAPES | 0.008 | 0.014 | 0.013 | 0.008 | -0.004 | 0.012 | -0.01 | -0.019 | -0.098 | -0.044 | -0.046 | -3.64 | -0.734 | -538566.554 |
| 0BK_BODY | 0.006 | -0.001 | 0.0 | -0.002 | -0.011 | -0.011 | -0.019 | -0.022 | -0.104 | -0.043 | -0.073 | -1.535 | -0.732 | -1245430.971 |
| 0BK_FACE | 0.005 | -0.004 | -0.003 | -0.006 | -0.011 | -0.017 | -0.018 | -0.025 | -0.098 | -0.077 | -0.1 | -1.599 | -0.711 | -617639.447 |
| STORY | 0.026 | 0.033 | 0.031 | 0.03 | 0.028 | -0.009 | 0.013 | -0.016 | -0.075 | 0.025 | -0.019 | -4.435 | -0.536 | -482269.634 |
| T | -0.016 | -0.018 | -0.017 | -0.018 | -0.025 | -0.018 | -0.025 | -0.02 | -0.113 | -0.085 | -0.092 | -1.383 | -0.898 | -236227.856 |
| FACES-SHAPES | 0.017 | 0.011 | 0.004 | 0.006 | -0.003 | -0.013 | -0.011 | -0.02 | -0.09 | -0.018 | -0.099 | -2.615 | -0.659 | -247146.447 |
| 2BK-0BK | 0.017 | 0.01 | 0.008 | 0.008 | 0.002 | 0.015 | -0.01 | -0.019 | -0.105 | 0.032 | -0.019 | -1.815 | -0.668 | -100109.264 |
| AVG | 0.011 | 0.006 | 0.007 | 0.007 | -0.003 | -0.001 | -0.006 | -0.016 | -0.092 | -0.037 | -0.076 | -1.774 | -0.718 | -194630.22 |
| T-AVG | -0.028 | -0.03 | -0.029 | -0.029 | -0.039 | -0.026 | -0.036 | -0.023 | -0.121 | -0.139 | -0.104 | -1.13 | -0.984 | -86920.081 |
| RF | -0.002 | -0.005 | -0.006 | -0.005 | -0.016 | -0.01 | -0.016 | -0.017 | -0.091 | -0.059 | -0.093 | -2.119 | -0.728 | -179995.523 |
| LH | -0.005 | -0.006 | -0.007 | -0.005 | -0.019 | -0.011 | -0.018 | -0.016 | -0.092 | -0.074 | -0.078 | -2.346 | -0.709 | -63500.533 |
| LH-AVG | -0.022 | -0.02 | -0.023 | -0.021 | -0.032 | -0.024 | -0.034 | -0.018 | -0.091 | -0.102 | -0.092 | -3.225 | -0.826 | -499450.47 |
| RANDOM-TOM | 0.004 | -0.001 | -0.001 | -0.002 | -0.009 | 0.007 | -0.015 | -0.015 | -0.084 | -0.004 | -0.055 | -4.323 | -0.683 | -18761.948 |
| LF | -0.003 | -0.005 | -0.007 | -0.005 | -0.015 | -0.013 | -0.016 | -0.017 | -0.092 | -0.058 | -0.093 | -2.071 | -0.769 | -331533.112 |
| PLACE-AVG | -0.007 | -0.017 | -0.017 | -0.016 | -0.02 | -0.002 | -0.025 | -0.02 | -0.113 | -0.048 | -0.133 | -1.846 | -0.805 | -15150.453 |
| MATH | 0.013 | 0.009 | 0.009 | 0.011 | 0.005 | -0.02 | -0.009 | -0.018 | -0.083 | 0.01 | -0.035 | -4.228 | -0.593 | -242997.845 |
| RH | -0.006 | -0.008 | -0.007 | -0.008 | -0.018 | -0.009 | -0.017 | -0.017 | -0.091 | -0.062 | -0.075 | -1.98 | -0.763 | -49992.747 |
| FACE-AVG | -0.002 | -0.013 | -0.014 | -0.015 | -0.02 | -0.019 | -0.025 | -0.023 | -0.112 | -0.028 | -0.116 | -1.482 | -0.818 | -210027.135 |
| RH-AVG | -0.026 | -0.025 | -0.025 | -0.026 | -0.033 | -0.025 | -0.032 | -0.02 | -0.101 | -0.096 | -0.081 | -2.147 | -0.828 | -44493.434 |
| RF-AVG | -0.027 | -0.025 | -0.026 | -0.027 | -0.033 | -0.024 | -0.031 | -0.017 | -0.096 | -0.096 | -0.12 | -2.259 | -0.819 | -27104.002 |
| LF-AVG | -0.026 | -0.028 | -0.03 | -0.028 | -0.034 | -0.025 | -0.033 | -0.017 | -0.104 | -0.096 | -0.123 | -2.557 | -0.858 | -217395.485 |
| BODY-AVG | -0.022 | -0.026 | -0.025 | -0.025 | -0.033 | -0.026 | -0.032 | -0.02 | -0.109 | -0.043 | -0.096 | -1.575 | -0.853 | -91476.217 |
| MATCH-REL | -0.02 | -0.017 | -0.017 | -0.015 | -0.022 | -0.008 | -0.018 | -0.021 | -0.099 | -0.032 | -0.051 | -3.646 | -0.752 | -20170.165 |
| TOOL-AVG | -0.026 | -0.032 | -0.032 | -0.031 | -0.04 | -0.034 | -0.042 | -0.021 | -0.111 | -0.033 | -0.062 | -1.173 | -0.836 | -20230.362 |
| PUNISH-REWARD | -0.036 | -0.036 | -0.035 | -0.036 | -0.038 | -0.03 | -0.037 | -0.02 | -0.114 | -0.034 | -0.047 | -1.622 | -0.949 | -75151.823 |

**Suppl. Table S2.** Weighted $R^2$ Score Table: $R^2$ Scores across all models examined ordered by Contrast names as in figure 4. These scores are provided in a CSV (model_r2w_scores.csv).

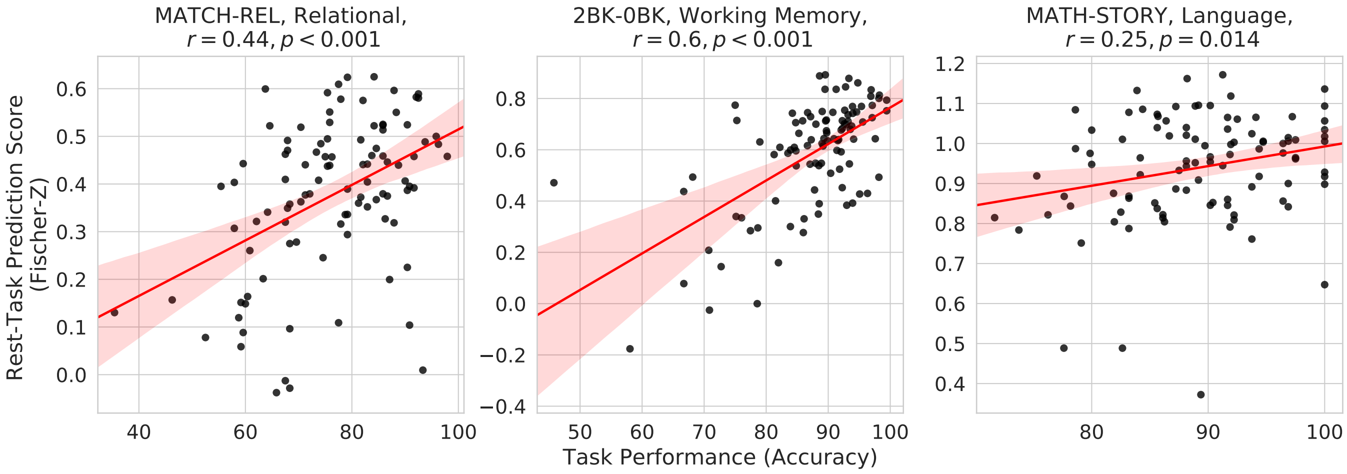


**Suppl. Figure S11.** Cortex-wise predictions scores of model **MMP-RR-PCR** plotted against behavior task accurary commited during the acquisition of the task. Three contrasts were chosen; Working Memory: 2BK-0BK, Relational: Match-Rel, and Language: Math-Story. Strong correlations are seen for all 3 contrasts.
